# Cancer-associated SF3B1 mutation K700E causes widespread changes in U2/branchpoint recognition without altering splicing

**DOI:** 10.1101/2024.11.18.624191

**Authors:** Andrey Damianov, Chia-Ho Lin, Jian Zhang, James L. Manley, Douglas L. Black

**Affiliations:** Department of Microbiology, Immunology, and Molecular Genetics, Molecular Biology Institute, David Geffen School of Medicine, UCLA Los Angeles, CA; Department of Biological Sciences, Columbia University, New York, NY

## Abstract

Myelodysplastic syndromes and other cancers are often associated with mutations in the U2 snRNP protein SF3B1. Common SF3B1 mutations, including K700E, disrupt SF3B1 interaction with the protein SUGP1 and induce aberrant activation of cryptic 3’ splice sites (ss), presumably resulting from aberrant U2/branch site (BS) recognition by the mutant spliceosome. Here, we apply the new method of U2 IP-seq to profile BS binding across the transcriptome of K562 leukemia cells carrying the *SF3B1* K700E mutation. For cryptic 3’ ss activated by K700E, we identify their associated BSs and show that they are indeed shifted from the WT sites. Unexpectedly, we also identify thousands of additional changes in BS binding in the mutant cells that do not alter 3’ ss choice. These new BS are usually very close to the natural sites, occur upstream or downstream, and either exhibit stronger base-pairing potential with U2 snRNA or are adjacent to stronger polypyrimidine tracts than the WT sites. The widespread imprecision in BS recognition induced by K700E with limited changes in 3’ ss selection supports a positive role for SUGP1 in early BS choice and expands the physiological consequences of this oncogenic mutation.

## Introduction

Pre-mRNA splicing is an essential step in gene expression and mutations in general splicing factors cause many forms of human disease (1–3). These mutations are mostly missense, usually cause slight alteration of function, and when assessed across the transcriptome may induce only modest changes in splicing. The core spliceosomal protein SF3B1 is commonly mutated in cancers (1, 4), with recurrent point mutations affecting residues of its HEAT repeat domains H4–H12 (1, 4–7). *SF3B1* mutations are found in approximately 30% of patients with myelodysplastic syndrome (MDS) (1, 3), with one mutation, K700E, accounting for more than 50% of these (5, 8). K700E and other *SF3B1* mutations, which are always heterozygous, induce recognition of cryptic 3′ splice sites (ss) in a small fraction of introns across the transcriptome. These new 3’ ss are often located upstream of the wild-type (WT) 3’ ss and are associated with an alternative upstream branch site (BS) (7, 9–11).

Recognition of the BS occurs in several steps during early spliceosome assembly. Initially, the proteins SF1, U2AF2 and U2AF1 recognize the BS, the polypyrimidine tract and the AG dinucleotide of the 3’ ss, respectively. These early factors recruit the U2 snRNP (U2), which after structural rearrangements forms the pre-spliceosomal A-complex with the BS base-paired to the U2 snRNA, leaving the future branchpoint (BP) adenosine bulged from the U2/BS helix (12–15). In both the A complex and an earlier complex prior to BS pairing, SF3B1 is in contact with U2AF and with intron sequences flanking the BS, thereby stabilizing U2 binding to the BS (16). There is substantial interest in how mutations in SF3B1 alter BS recognition, both for their relevance to cancer and for gaining insight into SF3B1 function within the spliceosome.

The K700E mutation has been shown to disrupt the interaction of SF3B1 with the splicing factor SUGP1, whose depletion or mutation recapitulates to a significant degree the splicing changes observed in SF3B1-K700E cells (10, 17, 18). SUGP1 is a G patch-containing protein that can bind to and activate DHX15, a DEAH-box RNA helicase proposed to function in several steps of spliceosome assembly and disassembly (19–22). Supporting the significance of the SUGP1-DHX15 interaction with respect to mutant SF3B1 splicing defects, it was found that a DHX15-SUGP1 G-patch fusion protein could rescue several aberrant splicing events (19). SUGP1 is thought to act early in BS selection where it contacts SF3B1, as well as SF1 and/or U2AF2. By one model, SUGP1 could then stabilize prespliceosomes prior to BS recognition by activating DHX15 to displace SF1, thereby allowing the future BS to base-pair with U2 snRNA. In the absence of SUGP1, SF1 may not be efficiently displaced diverting assembly onto a cryptic BS if available (23).

Other proteins affecting U2 assembly onto the pre-mRNA have also been implicated in aberrant BS selection by SF3B1 mutants. The G-patch protein GPATCH8 was also shown to modulate the activity of DHX15, and to functionally oppose the action of SUGP1 on approximately 30% of the splicing events altered in SF3B1-K700E mutant cells. This was suggested to reflect a competition between SUGP1 and GPATCH8 for DHX15, such that upon GPATCH8 depletion the excess free DHX15 could associate with mutant SF3B1 and partially rescue its splicing defects (24). Two other RNA helicases, the DEAD-box proteins DDX42 and DDX46 that play roles in initial assembly of the U2 on the pre-mRNA, were also proposed to affect BS selection by mutant SF3B1, although this has not been established (25–27). Thus, it is likely that K700E and other SF3B1 mutations affect the interactions of multiple spliceosomal proteins across the early steps of spliceosome assembly.

It has so far been difficult to directly assess BS use in the presence of *SF3B1* mutations (7, 11). Mammalian BS’s are highly degenerate and vary in their distance from the 3’ ss, making them difficult to predict with certainty. Introns can also have multiple BPs that are used at different frequencies. Large scale approaches to BP identification have used sequencing of the lariat intermediates or products of splicing, or applied machine-learning algorithms trained on large lariat sequencing datasets (28–32). These approaches have identified hundreds of thousands of BPs in human cells and tissues; however, they lack the capacity to quantify differential BS use between samples (29).

We recently developed U2 IP-seq as a whole-transcriptome method for isolating and sequencing BS’s base-paired with U2 snRNA in the precatalytic spliceosome (33). This method yields many unique BS RNA fragments per intron whose mapping and quantification allows comparison of BS use across different samples. To examine comprehensively how an SF3B1 mutation alters BS selection, we applied IP-seq to map sites bound by U2 in K562 cells expressing SF3B1 K700E and compared these maps to those for WT SF3B1. We find that the K700E mutation leads to far more widespread changes in U2/BS interactions than previously anticipated, with a large majority of these changes not affecting 3’ ss choice. By comparing sites showing differential U2 binding, we identify features that distinguish BSs favored by mutant SF3B1. Our findings provide new insights into both the normal functioning of the spliceosome and how mutations in SF3B1 alter this function.

## Results

### U2 IP-seq identifies numerous shifts in branch site recognition in MDS mutant cells

To obtain pre-mRNA-bound U2 complexes containing WT or mutant SF3B1, we isolated nuclei from two CRISPR-engineered sublines of K562 myelogenous leukemia cells. In each of these cell lines, one of the *SF3B1* alleles was gene-edited with either the WT sequence carrying synonymous mutations or with the K700E sequence, each fused to an N-terminal His6-FLAG epitope tag (10). Following the protocol previously described for HEK293 cells, the high-molecular weight (HMW) nuclear fraction containing chromatin was separated from the soluble nucleoplasm, and chromatin-bound complexes were extracted with a combination of DNAse and RNAse (33). The nucleoplasmic fraction was treated in parallel. We immunopurified U2 complexes from both fractions with either anti-Flag or an antibody binding a C-terminal peptide of SF3A2 (Figures 1, S1). The protein profiles of complexes purified from the HMW fraction of WT or SF3B1-K700E cells were similar (Figures 1, S1, right), and resembled those previously isolated from 293Flp-in cells (33). As seen previously, the SF3A and SF3B subunits were both abundant in the complexes isolated from the HMW fraction, as were the U2 snRNP proteins DHX15, and RBM17, while the Sm core and U2 A’/B” proteins were lost after nuclease digestion. PRPF8 and some other U5 components of the tri-snRNP were observed at lower stoichiometry in the immunopurified complexes. We did not detect SUGP1 protein in any of the complexes (Figure S1). Interestingly, immunopurified complexes isolated from the nucleoplasm contained only SF3A or SF3B subunits but not both. This may indicate that the nuclease treatment causes the disassociation of SF3A and SF3B from the U2 snRNP in this fraction, which is likely in the precatalytic 17S form or another complex unbound to pre-mRNA (34) (Figures 1, S1, left).

**Figure 1.**
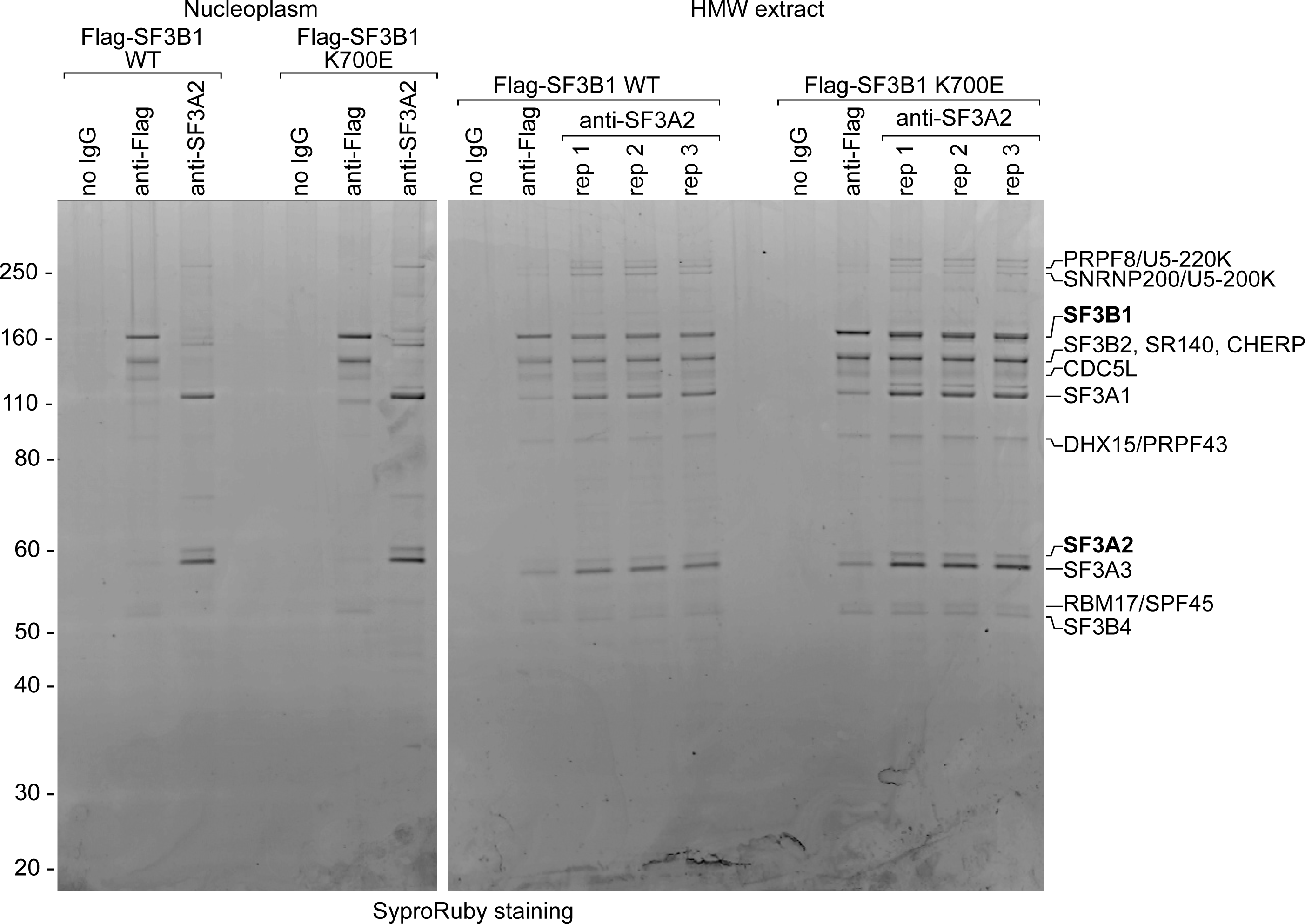
SF3B1 WT and K700E mutant proteins copurify with the same sets of U2 snRNP proteins in RNase-resistant complexes from the HMW nuclear fraction. Protein profiles of U2 complexes purified from soluble nucleoplasm (left) or HMW nuclear material (right) following extraction with RNase and Benzonase. Complexes containing SF3B1-Flag protein or SF3A2 were purified from WT and K700E cells with the indicated antibodies. Incubation with agarose beads without antibody served as negative control (no IgG lanes). The major coprecipitating proteins detected by protein staining are indicated on the right.

We isolated RNA from the HMW-extracted complexes and the U2 snRNA was removed from these samples by antisense oligonucleotide-directed RNase H degradation. The remaining population of protected RNA fragments were of similar size to those observed previously (Figure S2). These fragments were converted to cDNA, sequenced, and mapped to the human genome as described (33). As expected, the majority of the reads mapped in clusters near the 3’ ends of introns, corresponding to BSs protected by precatalytic spliceosomes (Table S1, Figure S3A). Putative BPs were scored within the clusters of BS reads according to match to consensus and position within the protected RNA fragment (33). IP-seq produces homogeneous peaks of reads at each BS with a low-background, and analysis of individual reads within each cluster allows identification of multiple BPs in many introns. The relative use of these alternative BPs was calculated as the ratio of reads supporting a particular BP to the total number of reads within the cluster. Detected BPs were graphically represented in browser tracks as vertical bars with the bar height proportional to this ratio (Figures 2, S3A-E, S4). IP-seq with the anti-SF3A2 antibody was performed in triplicate in both the WT and K700E cells and yielded highly reproducible results (Figures 2, S5A). For comparison, we also isolated U2 via the Flag-tag on SF3B1 from both WT and K700E cells. This yielded very similar peak profiles across the transcriptome (Figures 2, S3A).

**Figure 2.**
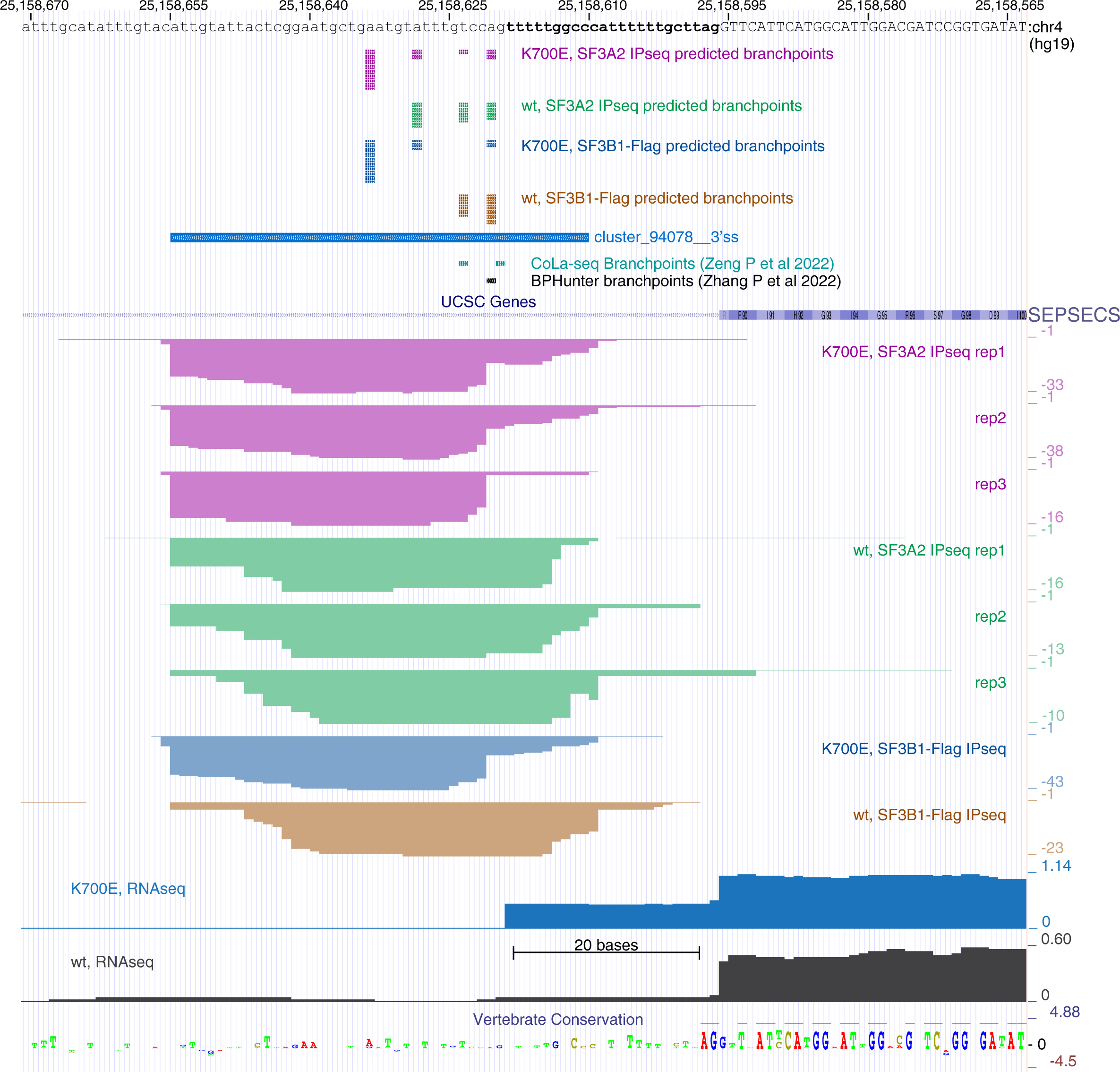
A branch site is selectively engaged with U2 snRNP in K562 SF3B1-K700E mutant cells, upstream of a cryptic 3’ splice site activated by this mutation. UCSC Genome browser view of the SEPSECS exon 4 and the upstream intron. Total read coverage is shown below the gene diagram. Individual IP-seq tracks show RNA recovered from the SF3B1-Flag WT or K700E mutant complexes, or from SF3A2-containing complexes from cells expressing the WT or mutant SF3B1 protein, each in triplicate. The significant branch site cluster of overlapping reads is marked in a track above the peaks, as are tracks of branch points identified by others (28, 31, 32). Precatalytic branch points predicted from individual IP-seq reads are shown above as vertical bars with the bar height proportional to the read numbers within the cluster as described in (33). Tracks of RNAseq data for the mutant and wildtype cells is at the bottom showing alternative 3’ splice site use. Other gene annotation and conservation tracks are shown as indicated.

Branchpoint positions across the transcriptome were found to be very similar between WT and SF3B1-K700E cells (Figure S5B). To specifically examine BPs upstream of differentially utilized 3’ splice sites (35), alternative splicing events were profiled using rMATS (36). This identified 339 alternative 3’ ss events that changed between WT and K700E (Table S2). To compare these events with the IP-seq data we removed events with low read coverage, or where the alternative 3’ ss co-occurred with other splicing patterns, such as exon skipping or intron retention. This produced a list of 157 events where both splicing choice and BP position could be confidently determined (Figures 3AB, S6, Table S3). For the majority of alternative 3’ ss events (143), we detected a shift in BP selection between the WT and K700E cells. Most of these splicing events involved the mutant protein shifting to an upstream BS and activating an upstream 3’ss (see examples with positive ΔPSI values in Figures 3A, S6, and Table S3). There were a minority of events (42) where the K700E protein shifted to a downstream BS and activated a downstream 3’ ss (negative ΔPSI values in Figures 3A, S6, and Table S3). For 3’ splice sites activated in mutant cells we did not find a notable difference in the average BP to AG dinucleotide distance (Figures 3A, S6). These results are consistent with previous RNAseq analyses of splicing changes induced by mutant SF3B1, where new splice sites used in mutant cells were usually upstream (7, 9–11). The distances between the WT and mutant BP ranged from a few to over 1000 nt, with the large majority less than 100 nt. The distances between the two 3’ss correlated strongly with these BP distances (Figure S6). This is predicted if the first AG downstream from the activated BP is chosen for splicing in the second catalytic step, as shown in earlier work (37, 38). The exception to this correlation was for BP to BP distances shorter than 20 nt, where the AG to AG distances remained clustered around 20 nt (Figure 3B and S6). This presumably reflects the exclusion of AG’s from the polypyrimidine tracts upstream from the WT AG. In these cases, the new AG is often within the protected fragment of the WT SF3B1 (Figure 2 and S3C).

**Figure 3.**
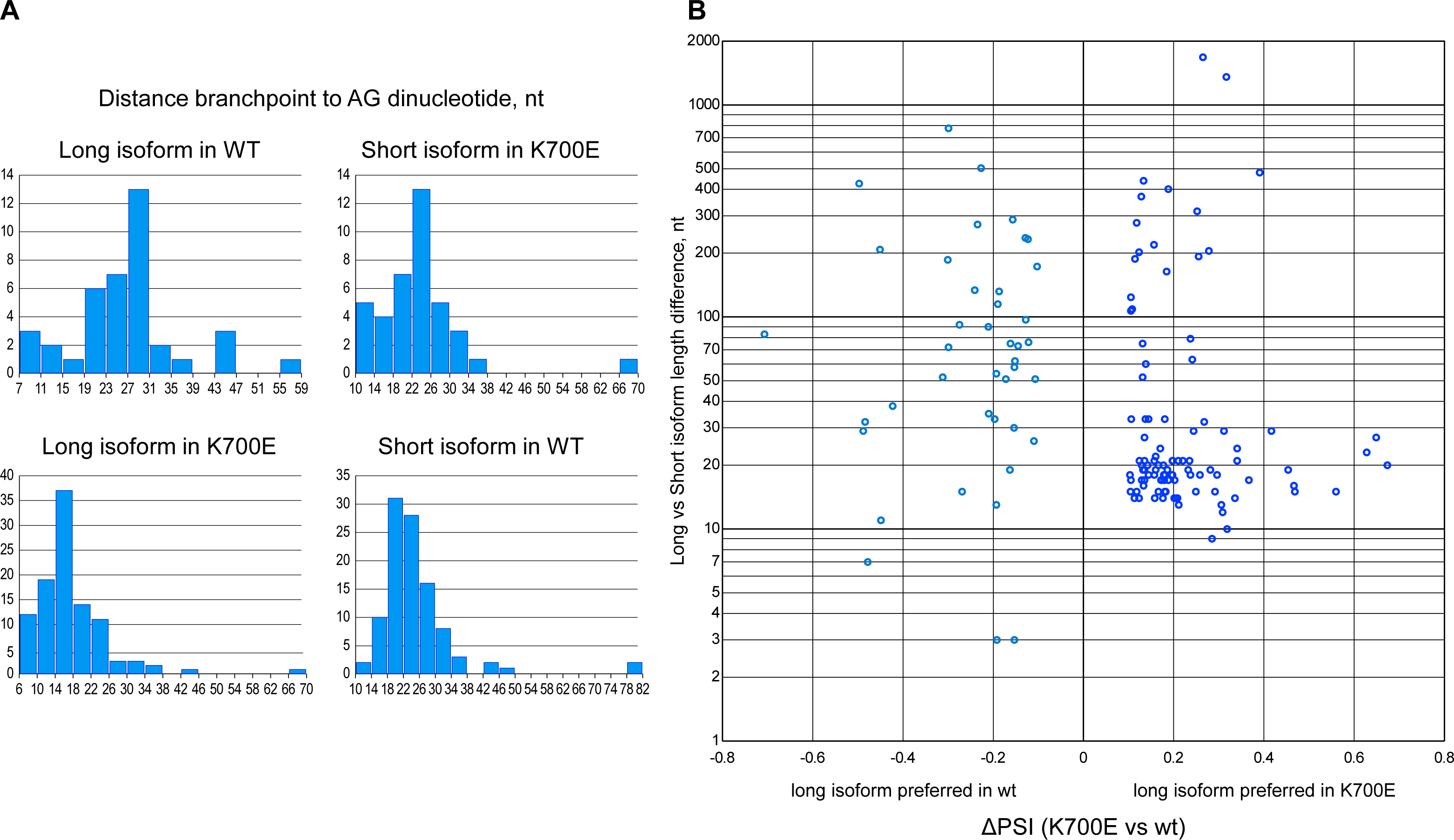
Distances between 3’ splice sites and associated branchpoints differentially used in K562 SF3B1 WT and K700E mutant cells (Related to Figure 2). **A.** Distribution of the nucleotide distances between the preferred branchpoint and the AG dinucleotide at the end of the intron is plotted for each short and long isoform resulting from alternative 3’ splice site use. **B.** Isoform length differences are plotted against the long isoform inclusion levels expressed in ΔPSI (percent spliced-in).

We also identified 14 alternative 3’ ss events where no alteration of BP use was detected despite the splicing shift (Table S3, example shown in Figure S4). However, examination of the BS clusters for some of these events indicated that reads for alternative BPs were present but were not sufficiently abundant to be reported as significant in our BP calling protocol (33).

It should be noted that in the cells expressing the mutant SF3B1-K700E allele, the SF3A2 antibody will isolate U2 complexes containing the products of both the mutant and WT *SF3B1* alleles. In contrast, IP-seq with the anti-Flag antibody will isolate only BSs bound by either the monoallelically tagged mutant or WT SF3B1 depending on the gene-edited cell line used. When the new BP was sufficiently distant to generate a new read cluster, the anti-Flag IP-seq usually generated a taller peak (Figure S3B,D). However, even when K700E BP reads overlapped the WT reads within the same cluster, the new predicted BPs were called in both the anti-Flag and anti-SF3A2 data (Figures 2, S3C,E). The identification of the K700E-dependent BPs by SF2A2 IP-seq highlights the sensitivity of this method. MDS mutations are always heterozygous. The ability of the SF3A2 antibody to detect allele-specific BPs bound by endogenous U2 indicates that the method should allow assays in heterozygous cell systems without epitope tagging of U2 components.

### The K700E mutation induces thousands of shifts in branchpoints without a change in 3’ splice site

Most alternative BPs found in SF3B1-K700E cells have been identified through the analysis of RNAseq data for changes in 3’ ss. We next examined the IP-seq data for BP changes that occur without altering ss choice. To search for such sites, we applied stringent criteria to create comparison sets of WT and K700E BPs. We identified the most frequently used BP in each IP-seq cluster of the K700E mutant cells. We then removed clusters where the most frequent BP in the mutant cells coincided with the most frequent BP in WT cells. For the remaining clusters, if a BP was present in WT cells at the same position as the most frequent one in mutant cells, we required that its frequency be less than half that seen in SF3B1-K700E. Strikingly, this identified nearly four thousand BPs whose use in mutant cells was greater than in WT cells (Table S4). Within each intron, we measured the nucleotide distance from the preferred mutant BP to that preferred in WT. Notably, these BP pairs were more narrowly spaced than those discovered by their proximity to alternative 3’ splice sites (compare Figure 4 and Table S3 column R). The majority of these mutant-preferred BPs were very close to the major BP used in WT cells (1-4 nucleotides) (Figure 4). At this distance, mutant preferred sites were most frequently but not exclusively found downstream of the WT preferred sites.

**Figure 4.**
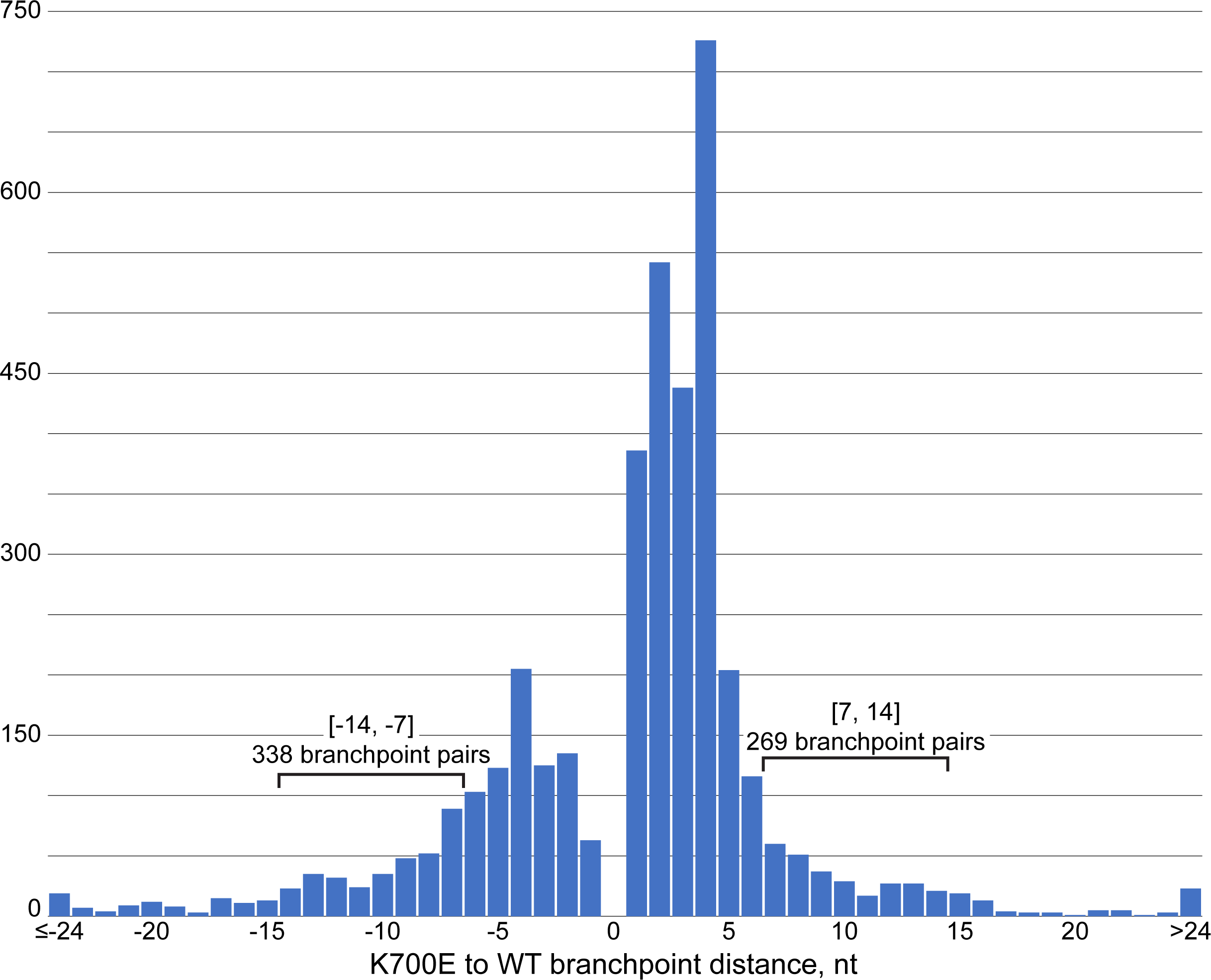
Branch sites selected in SF3B1-K700E mutant cells can be upstream or downstream of the site used in WT cells. Distribution of distances between branchpoints preferred in K562 SF3B1 mutant cells and those preferred in WT. Negative distances indicate mutant sites located upstream of WT, positive distances indicate mutant sites located downstream of WT. Only branchpoints located within the same branch site cluster are analyzed. The ranges of distances for branch site pairs selected for downstream analyses are indicated by brackets.

With this short separation, the nucleotide regions pairing with U2 snRNA at each site partially overlap, limiting the allowable nucleotide sequences between the two sites. To identify features of BSs preferred in SF3B1-K700E cells, we focused on those sufficiently distant from the preferred WT BP that the base-paired segments did not overlap. We selected 338 BP pairs where the mutant preferred site was 7 to 14 nucleotides upstream of the WT branchpoint, and 269 pairs where the mutant preferred site was 7 to 14 nucleotides downstream of the WT (Figure 4).

The U2 snRNA sequence GUAGUA can bind BSs in several base-pairing modes that differ in the positions of individual base pairs and the number of bulged nucleotides (29). To assess the relative use of different pairing modes in K562 cells, we examined a subset of K562 BSs that were independently mapped in two previous studies (28, 32).

Comparing the base-pairing potentials of these sites in each of the described pairing modes, we found possible examples of all six modes but at different relative frequencies than observed previously (29) (Figure S7 and Table S5). In particular, there were many fewer BPs predicted to use a C nucleotide instead of A than reported in the previous study. The canonical BS that uses U2 snRNA pairing mode 1 was favored over the other modes in K562 cells (Figure S7). The BPs preferred in alternative 3’ splice sites induced by mutant SF3B1 had similar distributions of modes for BPs both upstream and downstream from the WT site (Figure S8).

BPs that were closely spaced favored particular combinations of pairing modes, presumably due to sequence constraints on the nucleotides separating them. For BP pairs separated by 4 nucleotides, the most frequent pairing mode combinations were 1-1, 4-1, and 5-1. However, the sites preferred by the mutant protein did not show a bias for a particular pairing mode. Similarly, the consensus sequences of these closely spaced BS pairs did not show strong differences between the upstream and downstream K700E sites (Figure S9).

### K700E SF3B1 favors branchpoints with more stable pairing to U2 snRNA or with stronger polypyrimidine tracts

We next assessed what intronic features, if any, might distinguish the BSs preferred by the mutant SF3B1. To compare the U2 snRNA base-pairing potential of WT and mutant BSs separated by 7-14 nucleotides, we grouped them by pairing mode (Figure S10) and generated consensus sequences. For canonical mode 1 sites, upstream sites preferred in K700E cells exhibited more potential base-paired nucleotides with U2 snRNA than the WT sites downstream (Figure 5A, upper row). Interestingly, the mutant preferred sites downstream from WT did not necessarily exhibit greater base pairing than the WT site. Instead, these mutant-preferred downstream sites were adjacent to more prominent U-rich polypyrimidine tracts (Figure 5A, lower row). Similar findings were made for the mutant and WT BS pairs associated with cryptic 3’ splice sites (Figure 5B). BSs assigned to other pairing modes often showed similar trends for the mutant sites, but these enrichments were based on fewer examples and were not as statistically significant (Figure S11).

**Figure 5.**
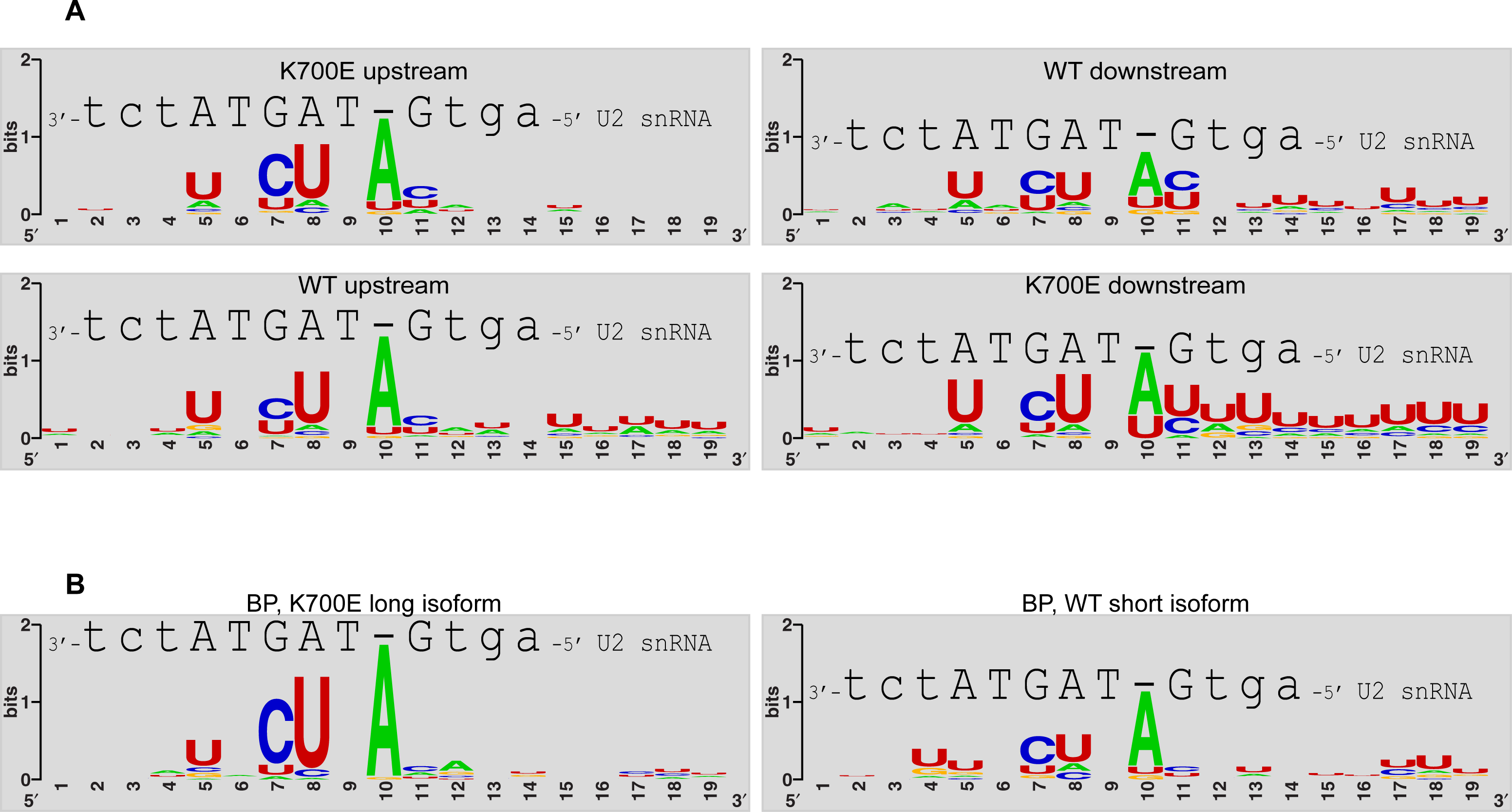
Upstream branch sites preferred in K700E mutant cells exhibited stronger matches to the consensus than the downstream WT site, whereas mutant-preferred downstream sites are adjacent to more U-rich polypyrimidine tracts. **A.** Consensus sequences derived from differentially used branch sites separated by 7 to 14 nt and their base-pairing with U2 snRNA in the canonical mode 1 are shown as in Figure S7B. **B.** Consensus sequences for mode 1 branch sites, associated with alternative 3’ splice sites. Only sites from introns where the longer isoform is produced in mutant cells are shown.

Overall, we have found that the *SF3B1* K700E mutation results in numerous previously undetected shifts to cryptic BSs by the mutant U2 snRNP. These widespread changes in BSs, which occur independently of changes in 3’ ss choice, are associated with the shift of U2 to more strongly interacting sites, either through enhanced base pairing with U2 snRNA or through a more U-rich polypyrimidine tract that may exhibit stronger U2AF2 binding.

## Discussion

We previously isolated U2 particles from the chromatin fraction of 293Flp-In cells and showed that they copurified with protected intron BSs across the transcriptome. We now show that U2 particles can also be isolated from K562 cells carrying either *SF3B1* WT or the K700E allele found in MDS and other cancers. U2 complexes with WT or mutant SF3B1 contained the same set of core U2 proteins, and were again stably bound to intron BSs across the transcribed RNA. The strong protection of intron BPs by the U2 particles indicates the formation of the U2/BS helix, while the lack of branch formation indicates that the particles are likely derived from A- or B-complex spliceosomes. Characterizing the RNA fragments protected from nuclease in these particles allowed us to compare U2/BS interactions in WT and mutant cells.

Previous studies demonstrated that multiple *SF3B1* mutations, including K700E, destabilize SF3B1 interaction with the protein SUGP1 and reduce its ability to activate the helicase DHX15 (8, 14, 20). SUGP1 has not been observed in the early stage spliceosomes so far characterized, and likely acts transiently prior to BS recognition (23). These data are consistent with a model in which SUGP1 mediates interactions between U2, SF1, and/or U2AF2 at the BS/3’ SS. The G patch of SUGP1 then activates DHX15 ATPase activity to displace SF1 from the pre-mRNA, allowing U2 to associate with the BS (23). In the absence of SUGP1, DHX15 is not activated, SF1 is not effectively removed from the BS, and U2 may shift to a cryptic BS if available. Notably, the U2 particles described here have already undergone BP assembly and lack SUGP1 (33). The observed binding of U2 to alternative BSs in the SF3B1 mutant cells indicates that assembly onto these sites is stable after the early SUGP1 step and persists into mature spliceosomes. The remarkable finding from our data is that the mutant spliceosome recognizes new BSs not just associated with the several hundred cryptic 3’ splice sites, but also binds to thousands of cryptic BSs that do not lead to alternative 3’ ss usage. It is likely that reducing the stringency of our search criteria would identify additional cryptic BS in the mutant cells that are used at lower frequency. Note that of the ∼124,000 mapped intronic BS clusters, only ∼4,000 showed changes in BP choice (∼3.2%). Although there are extensive changes in BP’s in the mutant cells, the majority of splicing across the transcriptome is occurring normally.

An alternative model for how loss of SUGP1 and the consequent failure to activate DHX15 leads to cryptic BS/3’ ss usage posits a quality control function whereby SUGP1/DHX15 induces prespliceosome disassembly when an erroneous BS is bound (20, 22). This model is consistent with another known role of DHX15, or its yeast homologue Prp43, which is to induce disassembly of the intron lariat complex in the final step of the splicing cycle (39–41). From our data, such an activity in the early spliceosome would require that WT U2 snRNP naturally recognize the thousands of cryptic BSs that we find in close juxtaposition to the authentic BS. This would in some way induce recruitment of SUGP1 and activation of DHX15 to selectively disassemble the cryptic BS complexes. As a seemingly more parsimonious explanation for the data, we favor the idea that SUGP1/DHX15 facilitates recognition of the correct BS rather than preventing use of incorrect BSs, although neither model is ruled out. Our finding that the majority of cryptic BSs used in the mutant cells provide a better match to the consensus than the WT site also seems consistent with a positive role for SUGP1/DHX15 in facilitating recognition of authentic sometimes weaker BSs. It is not clear in either model what features of a BS might allow SUGP1 to determine its use. We need to learn more about the very earliest steps of U2 snRNP assembly to understand fully the role of SUGP1/DHX15 in BP selection.

The distribution and number of reads recovered from a particular BS is consistent across samples (33), allowing identification of changes in BS preferences between cell populations. This was validated by the detection of differentially engaged BSs for a large fraction of the cryptic 3’ splice sites identified by RNAseq as activated by SF3B1-K700E (35). It is notable that many of these alternative 3’ splice sites were associated with multiple upstream BPs. In some cases, a new site not seen in WT was strongly preferred in mutant cells (Figure 2). In other cases, most or all the BSs could be observed in both WT and mutant cells, but their distribution changed. Although changes in later stages may occur, our finding of new BPs for nearly all alternative 3’ ss activated by K700E indicates that changes in spliceosome assembly have already occurred prior to the first catalytic step.

As noted above, the most unexpected finding from our data is that the new BSs bound by the mutant spliceosome are not limited to the those associated with a few hundred cryptic 3’ splice sites but include thousands of cryptic BSs not associated with alternative 3’ ss usage. It is well established that the same 3’ ss can be chosen in the second splicing step after branching to multiple BPs, and that the relative usage of these BPs adjacent to a single 3’ ss can change under different conditions (42). The use of novel BSs in MDS mutant cells that are not associated with changes in splicing has potentially important implications for the disease phenotype. For example, it is not understood whether alternate BPs might differ in performance during the later steps of spliceosome assembly and catalysis. BP changes could have complex effects on mRNA levels by altering the kinetics of splicing for certain introns, or the overall efficiency of product formation. Early in vitro studies indicated that BP choices can be altered by regulatory factors as well as have effects on the rate of splicing (42–44). Thus, the altered BS selection resulting from mutation of SF3B1 may affect cellular mRNA metabolism and gene function in more subtle ways than the identified changes in mRNA isoforms.

The BSs preferred in K700E mutant cells over WT showed several common features. Mutant preferred sites upstream of the WT BS exhibited stronger base pairing potential with U2 snRNA (Fig. 4). Similar observations were previously made for potential BSs associated with upstream 3’ splice sites activated by SF3B1 mutations at R625 and K666 (7), which also disrupt interaction with SUGP1 (8). In contrast, mutant preferred BSs downstream of WT sites were not markedly stronger in their U2 base-pairing potential than the WT sites but were instead adjacent to polypyrimidine tracts with greater enrichment for U residues. These findings fit with the model that the weakened SUGP1 binding to SF3B1 in the K700E cells may necessitate additional stabilization for U2 to engage a BS (23). This stabilization may come either from stronger U2 snRNA pairing or a more U-rich and more closely adjacent polypyrimidine tract, with higher U2AF2 occupancy (10, 45). A closer match to the BS consensus could also act to enhance SF1 binding (46). The branch sites preferred in K700E and WT cells were most frequently in close proximity and partially overlapping (Fig. 3). It may be that U2 snRNA can shift its pairing along the pre-mRNA to a more favorable interaction close by, or that different sites of SF1 binding can be favored prior to engagement of the BS by the incoming U2.

We did not find evidence for the mutant SF3B1 preferring particular modes of pairing for the U2 snRNA. Canonical pairing (Mode 1; (29)) was the most common in both WT and mutant cells and there were no clear preferences for a new pairing mode when the binding shifted. It is difficult to compare the relative stabilities of BS-U2 snRNA pairing in different modes, and examples of BSs using non-canonical pairing modes were more limited. Analyses of larger datasets combined with better assessment of the binding affinities for different RNA pairings may elucidate additional features affecting SF3B1 function in BS selection.

The K562 cells carrying the K700E mutation provide a valuable means for determining how an MDS mutation affects both splicing and BS recognition. The MDS phenotype is manifested in hematopoietic stem and progenitor cells, and it may be that different BP alterations occur in these cells. It should thus be informative to perform U2 IP-seq in MDS patient tissue samples or cells derived from patient iPSCs. Our study demonstrates the feasibility of such studies in its sensitive detection of new BPs in cells carrying a heterozygous *SF3B1* allele, and in using a U2 antibody to assay endogenous protein without an epitope tag. It will also be interesting to assay changes in U2 assembly resulting from other mutations in SF3B1 and in cells of other tumor types carrying splicing factor mutations.

In summary, we identify transcriptome-wide alterations in branch site recognition caused by a common cancer-associated mutation in the U2 snRNP protein SF3B1. Our findings give new insight into the mechanisms controlling spliceosome assembly, and are important for understanding the role of splicing misregulation in neoplastic disease.

## Methods

### Cell lines

K562 cells were grown in Iscove’s Modified Dulbecco’s Medium (IMDM) supplemented with 10% FBS in a 5% CO2 incubator (10). SF3B1-K700E mutant cells were inoculated at twice the density of the WT cells and grown for 72 hours.

### Antibodies

Primary antibodies used for immunoblot assays: SUGP1 (Thermo Fisher Scientific, A304-675A). SF3A1, SF3B1, and U1-70K are described (47). SF3A2 antibody: Rabbits were immunized with the human SF3A2 C-terminal peptide C_MLRPPLPSEGPGNIP, conjugated to Keyhole Limpet Hemocyanin (KLH). Following three boosts, IgG clones cross-reacting with the SF3A2 peptide were obtained by single B-cell sequencing (GenScript). These were coupled to Protein G-agarose (Pierce), crosslinked using DMP, and further screened for specific immunoprecipitation and efficient elution of U2 snRNP with excess MLRPPLPSEGPGNIP (SF3A2-CT) peptide.

### RNP sample preparation

Cells were harvested and washed in Phosphate Buffered Saline. Nucleoplasm and HMW material were extracted as described (33). Extracts were incubated overnight at 4 °C with 5-7.5 μl packed-volume of either M2 FLAG agarose beads (Millipore-Sigma), anti-SF3A2 Protein G-agarose beads, or Protein G-agarose crosslinked in absence of IgG. Beads were then washed four times with wash buffer (20 mM HEPES-KOH pH 7.5, 150 mM NaCl, 1.5 mM MgCl_2_, and 0.05% Triton-X100), and eluted for two hours at 4°C in presence of 150 ng/μl of either 3xFLAG (Millipore-Sigma) or SF3A2-CT peptide. Proteins analyses were performed by staining with SYPRO Ruby (Thermo Fisher Scientific) and immunoblotting. Aliquots of RNA obtained following deproteinization were subjected to 5’ radiolabeling and separated by Urea-PAGE, or were sequenced using Illumina technology as described (33).

### IPseq and RNAseq data analysis

Removal of PCR duplicates, genome alignment, analysis of clustered reads, and branchpoint prediction were performed as described (33). Branchpoint use within a defined 3’ splice site cluster was expressed as the ratio of the number of reads supporting a particular branchpoint to the total number of reads in the cluster. Branchpoint preference was calculated as the ratio of branchpoint use in WT vs K700E mutant samples. All sequences shown were aligned to the human genome hg19.

Alternative splicing was analyzed by rMATS (36) and expressed as changes in percent-spliced-in values (ΔPSI). 3’ splice site events showing a splicing change between control K700E and WT cells were considered regulatory targets (|ΔPSI|>10 with FDR less than 0.05). Differentially used branchpoints at these alternative 3’ splice sites were selected manually. Only one pair of branchpoints was chosen, even though multiple WT or K700E branchpoints were observed at some of these events.

Differentially used branchpoints not necessarily associated with alternative 3’ splice sites were selected from 3’ splice site and intronic IPseq clusters. K700E rank1 branchpoints were preferred at least two-fold compared to WT and were selected along with the rank1 branchpoint used in WT. Branchpoint pairs supported by fewer than 25 SF3A2 IPseq K700E and WT reads combined and not ranked in SF3B1 IPseq data were excluded.

Branch site base pairing with the U2 sequence GUAGUA in each of the six described modes (48) was assessed by awarding four points per GC pair, 3 points per AU pair, and two points per GU pair. Mismatches were not issued a penalty. The pairing mode with highest score, calculated as the sum of points, was selected as the most likely for each branch site.

## Acknowledgments

We thank Luke Buerer and William Fairbrother (Brown University, RI) for sharing code and helpful discussion. This work was supported by awards from the Jonsson Comprehensive Cancer Center at UCLA to D.L.B., and by NIH grants R01GM127473 (A.D.), R21HG012624 (D.L.B.), R35GM136426 (D.L.B.), and R35GM118136 (J.L.M.).

Sequencing data generated in this study has been deposited at NCBI GEO under the accession number GEO: GSE280659 (https://www.ncbi.nlm.nih.gov/geo/query/acc.cgi?acc=GSE280659).

## Supplementary table legends

**Table S1.** Summary of IP-seq data.

The numbers of genome-aligned reads and defined read clusters are listed for each of the four experimental datasets as indicated. Numbers and percentages of these reads and clusters mapping to branch sites, 5’ splice sites, and to undefined genomic sites are also listed. The numbers of SF3A2 IP-seq reads shown are combined from three biological replicates.

**Table S2.** SF3B1 K700E-dependent splicing changes.

rMATS identified 3’ splice site changes between SF3B1 K700E and WT cells. Only significant events with |ΔPSI| > 10 and FDR < 0.05 are listed. For each splicing event, columns list gene name, exon coordinates, splice junction counts, average inclusion level for each condition, p value, FDR, and delta PSI.

**Table S3.** Branchpoints differentially engaged in K700E and WT cells, upstream of alternative 3’ splice sites.

Splicing analysis data as in Table S2 is merged with differentially engaged branch site sequences associated with the short and the long spliced isoforms. For each branch site, a 19nt-long sequence is shown with the branchpoint nucleotide in the middle. The putative U2-base pairing mode of each site is also listed. Normalized ratios indicate the number of reads supporting the corresponding branchpoint, divided by the total number of reads in the cluster. The ratio of these normalized ratios in WT vs K700E cells, indicating preferential branchpoint selection, is also shown. Events where the same branchpoint appears to be used in both WT and K700E cells are listed in a separate tab.

**Table S4.** Differentially engaged branchpoints not associated with altered 3’ splice site choice.

Differentially engaged branchpoints are listed similarly to Table S3. The columns indicating the distance between branchpoints and the preference ratio are highlighted in yellow.

**Table S5.** A list of branch sites discovered in two studies of K562 cells (28, 32) were assessed for optimal base-pairing with the branch-site recognition sequence in U2 snRNA. Sequences and potential base-pairing modes are listed as in Table S3.

---

**Figure S1.**
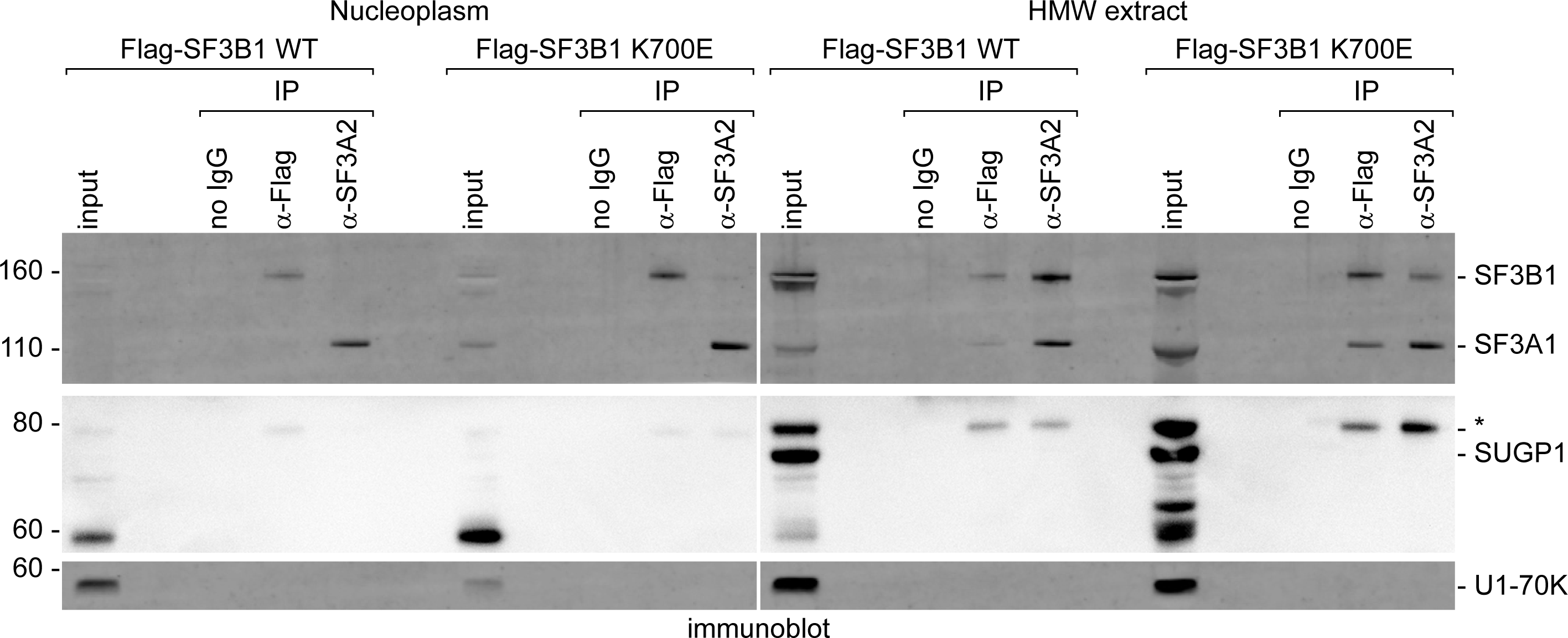
SF3B1 WT and K700E proteins do not coprecipitate detectable SUGP1 from nucleoplasmic or HMW extracts (Related to Figure 1). Complexes purified with SF3A2 and Flag antibodies in Figure 1 were subjected to immunostaining with SF3B1, SF3A1, and SUGP1 antibodies. Total nucleoplasmic and HMW extract corresponding to 5% input material was also probed. The protein band indicated by an asterisk is an unknown substoichiometric U2 snRNP associated protein stained with the SUGP1 antibody but migrating above SUGP1. Immunostaining with U1-70K IgG served as negative control.

**Figure S2.**
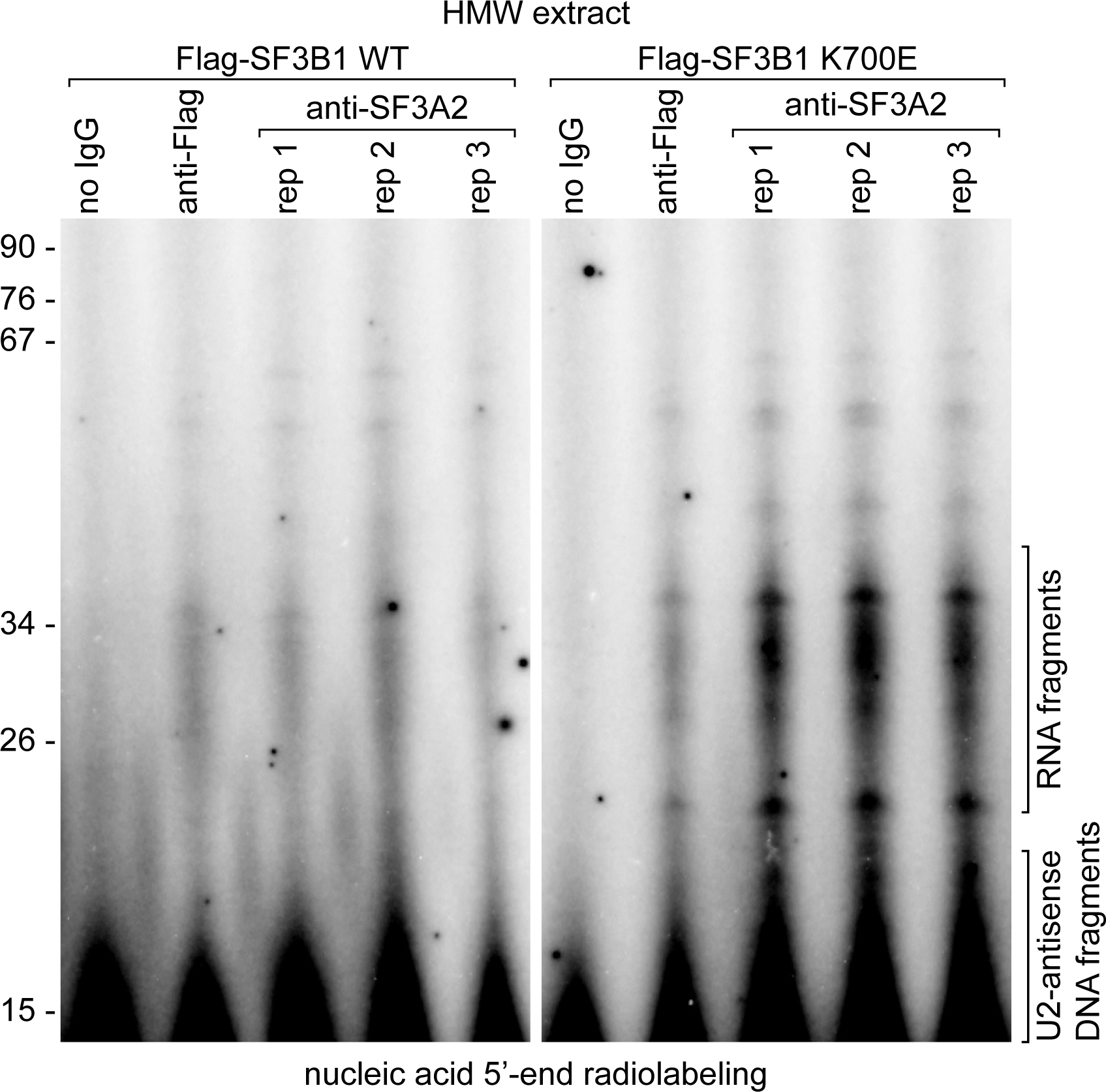
The SF3B1 WT and the K700E U2 snRNP complexes from the HMW fraction contain protected pre-mRNA (Related to Figure 1). 5’-radiolabeled RNA, recovered from SF3B1-Flag or SF3A2 immunopurified complexes shown in Figure 1 was separated by denaturing Urea-PAGE and detected by phosphorimaging. Samples were pretreated with full-length U2 snRNA-antisense DNA and RNase H, followed by TurboDNase digestion. Detected RNA fragments and short digested DNA oligonucleotides are indicated on the right. See Reference (33).

**Figure S3.**
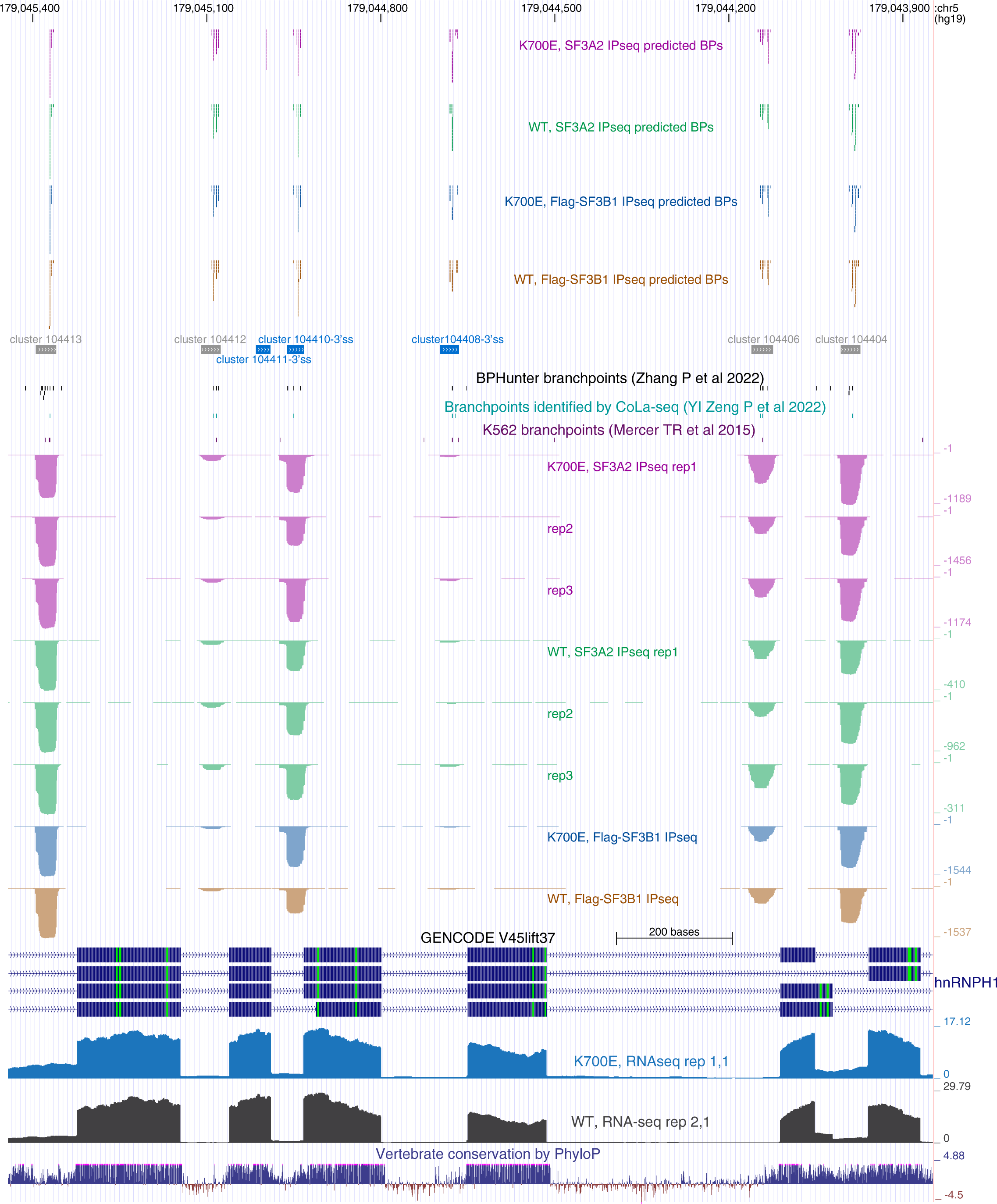
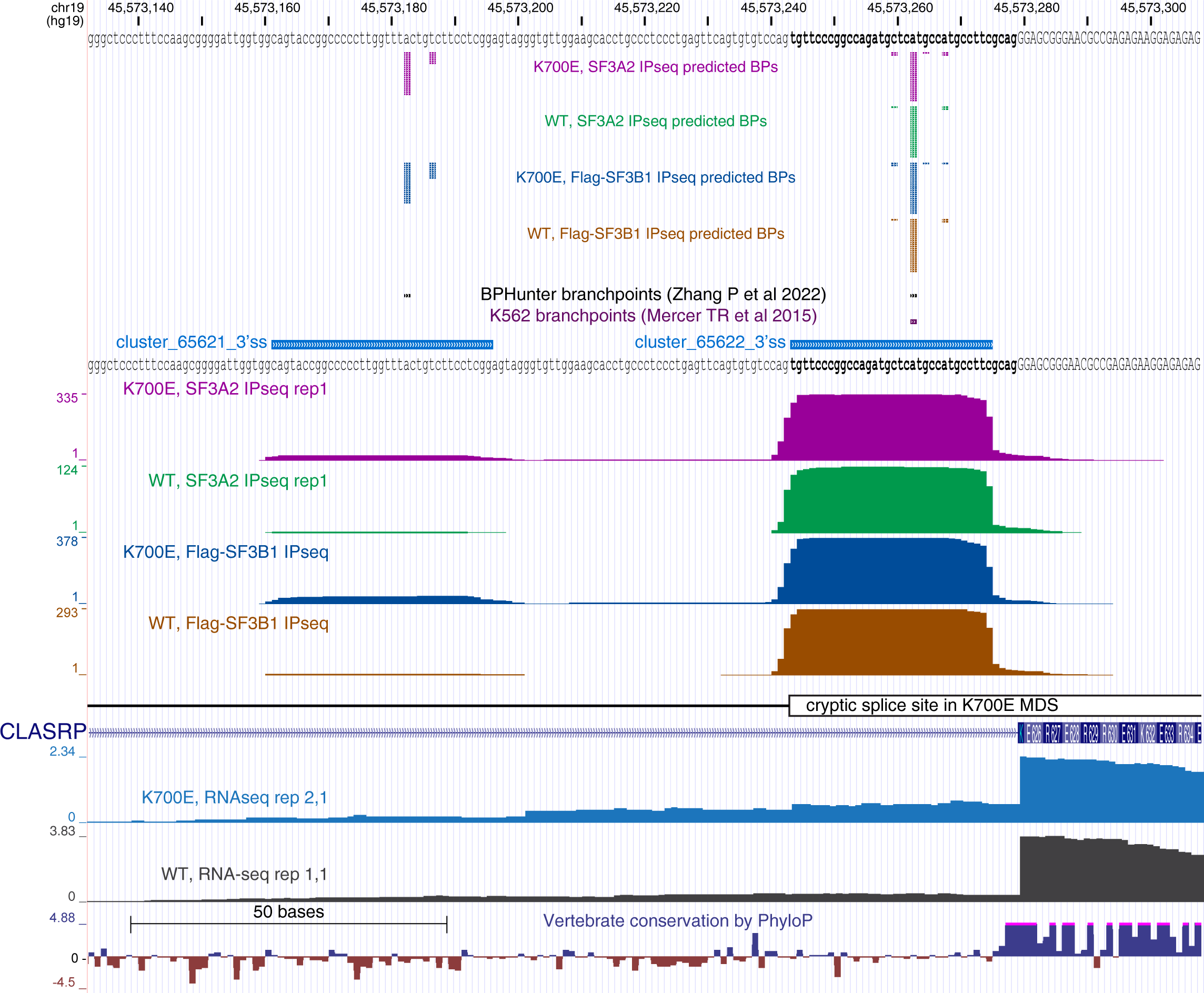
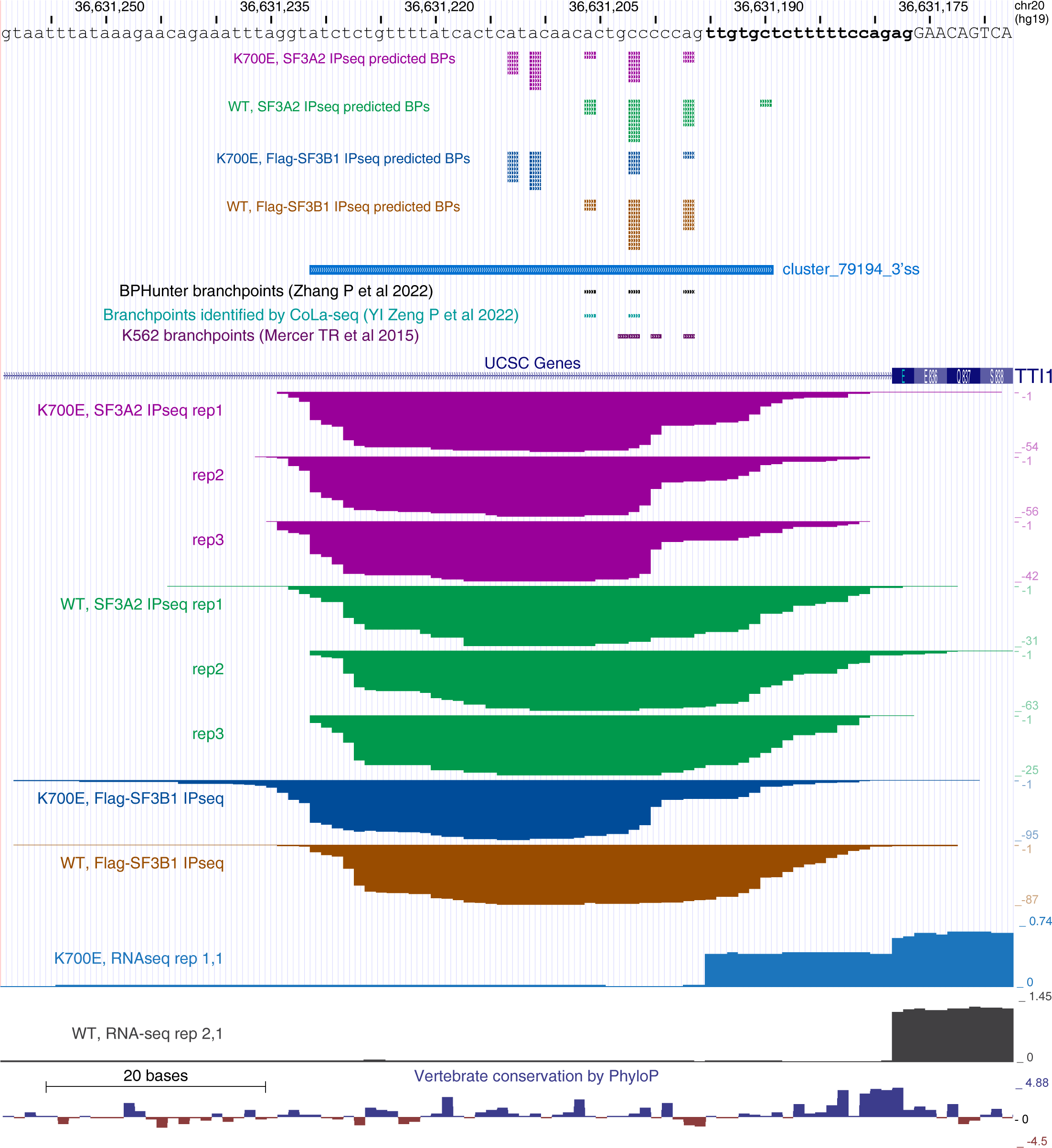
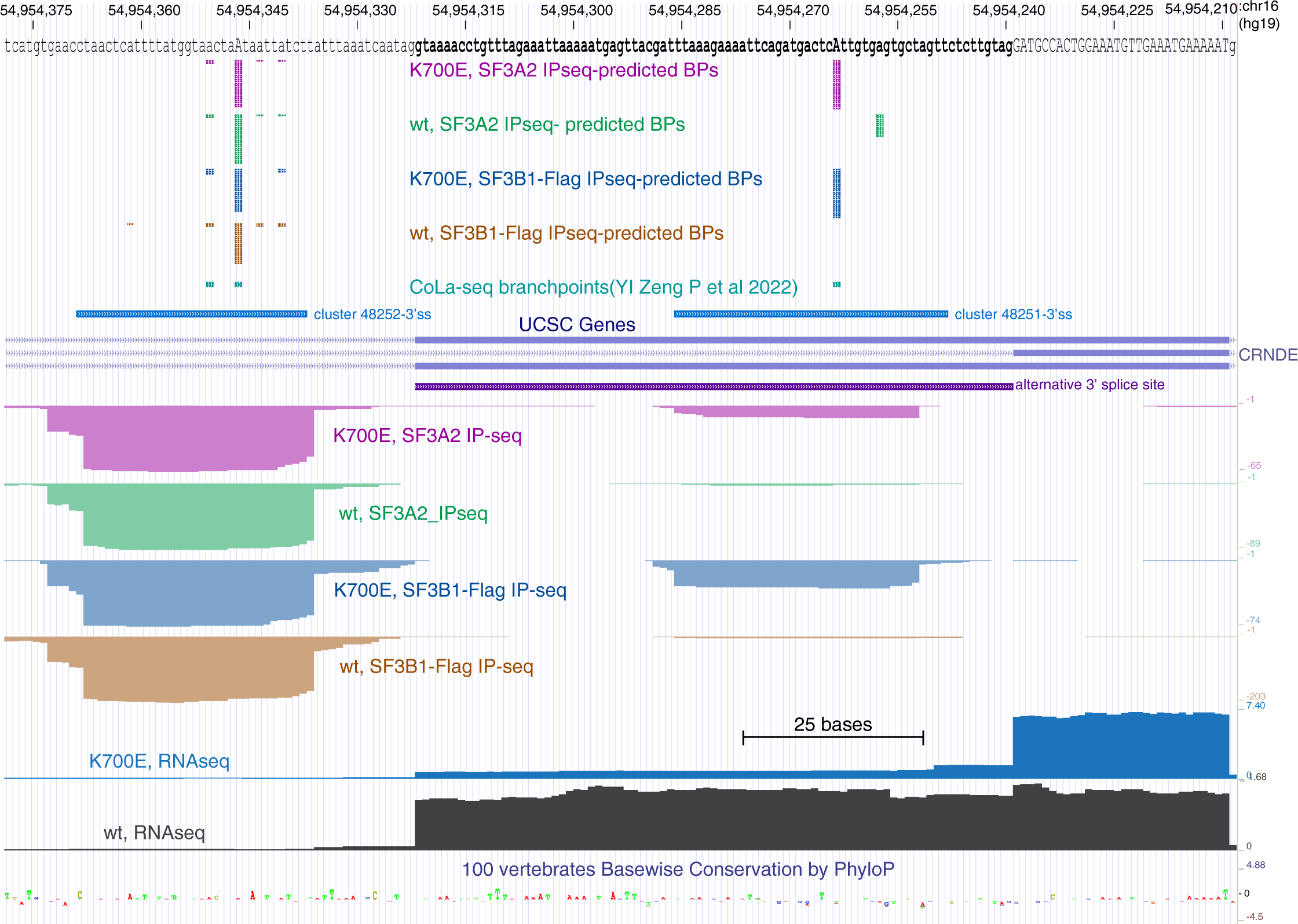
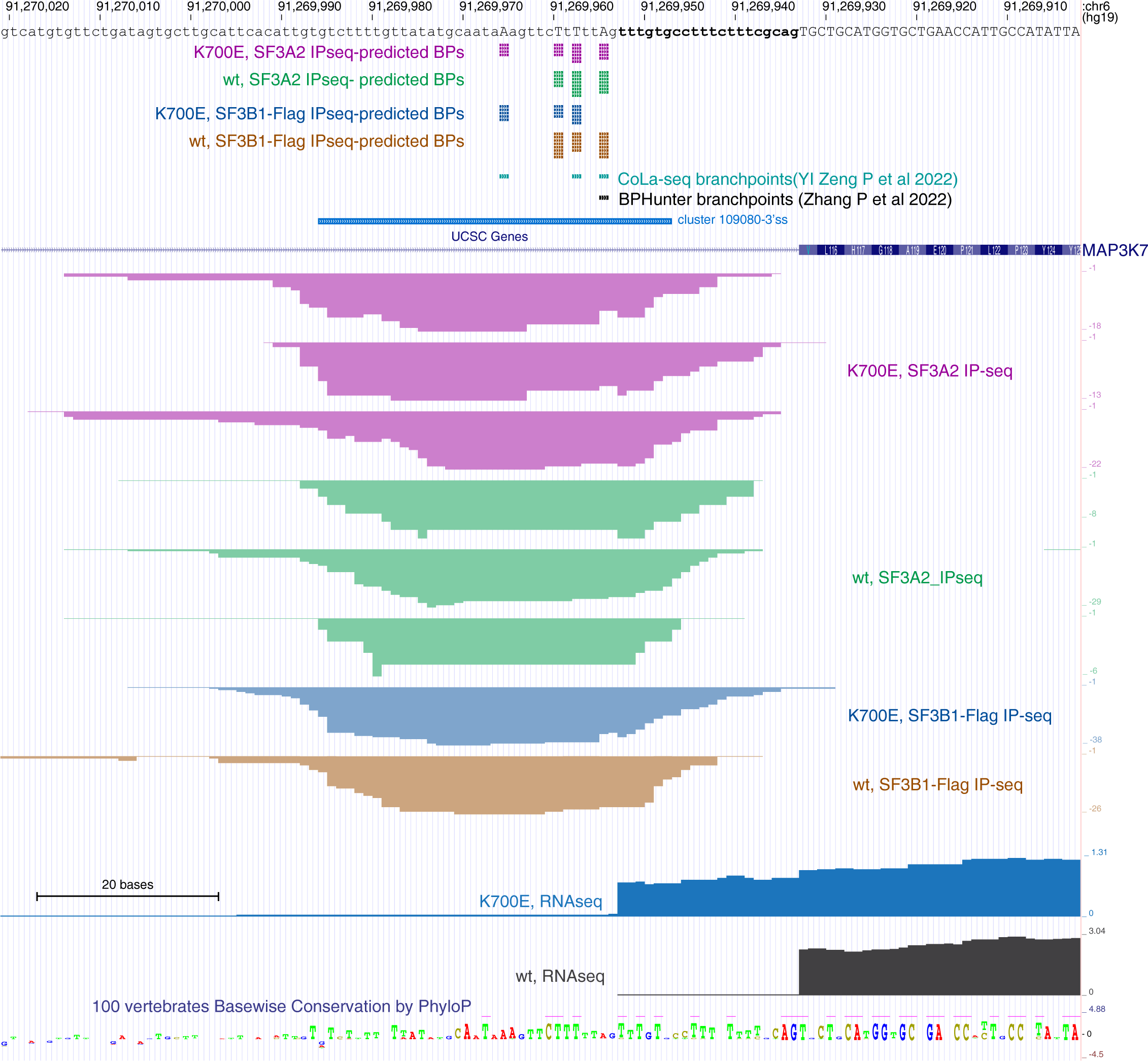
Examples of the RNA protection patterns for the SF3B1 WT and K700E mutant complexes bound to pre-mRNA branch sites (Related to Figure 2). **A.** UCSC Genome browser view of branch sites detected by IP-seq mapped to a portion of the hnRNPH1 gene. Branch sites detected at aberrant 3’ splice sites activated by SF3B1-K700E are shown for **B.** CLASPR, **C.** TTI1, **D.** CRNDE, and **E.** MAP3K7 genes. Genomic tracks are shown as in Figure 2.

**Figure S4.**
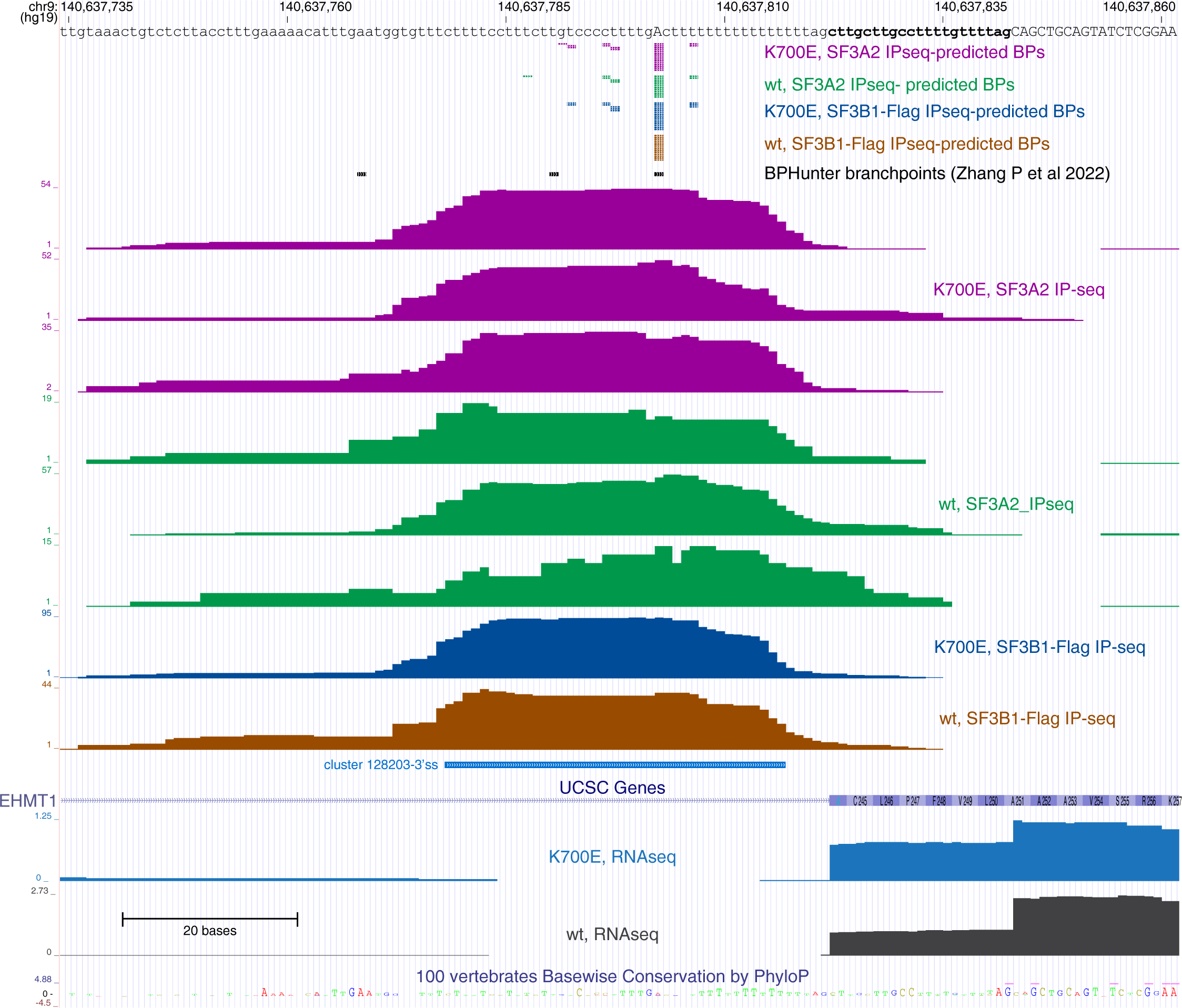
Alternative 3’ splice site upstream of EHMT1 exon 5 is engaged in K562 SF3B1-K700E mutant cells without detection of a different branch site. Predicted branchpoints, IP-seq, RNA-seq, and other genomic tracks are also shown as in Figure 2.

**Figure S5.**
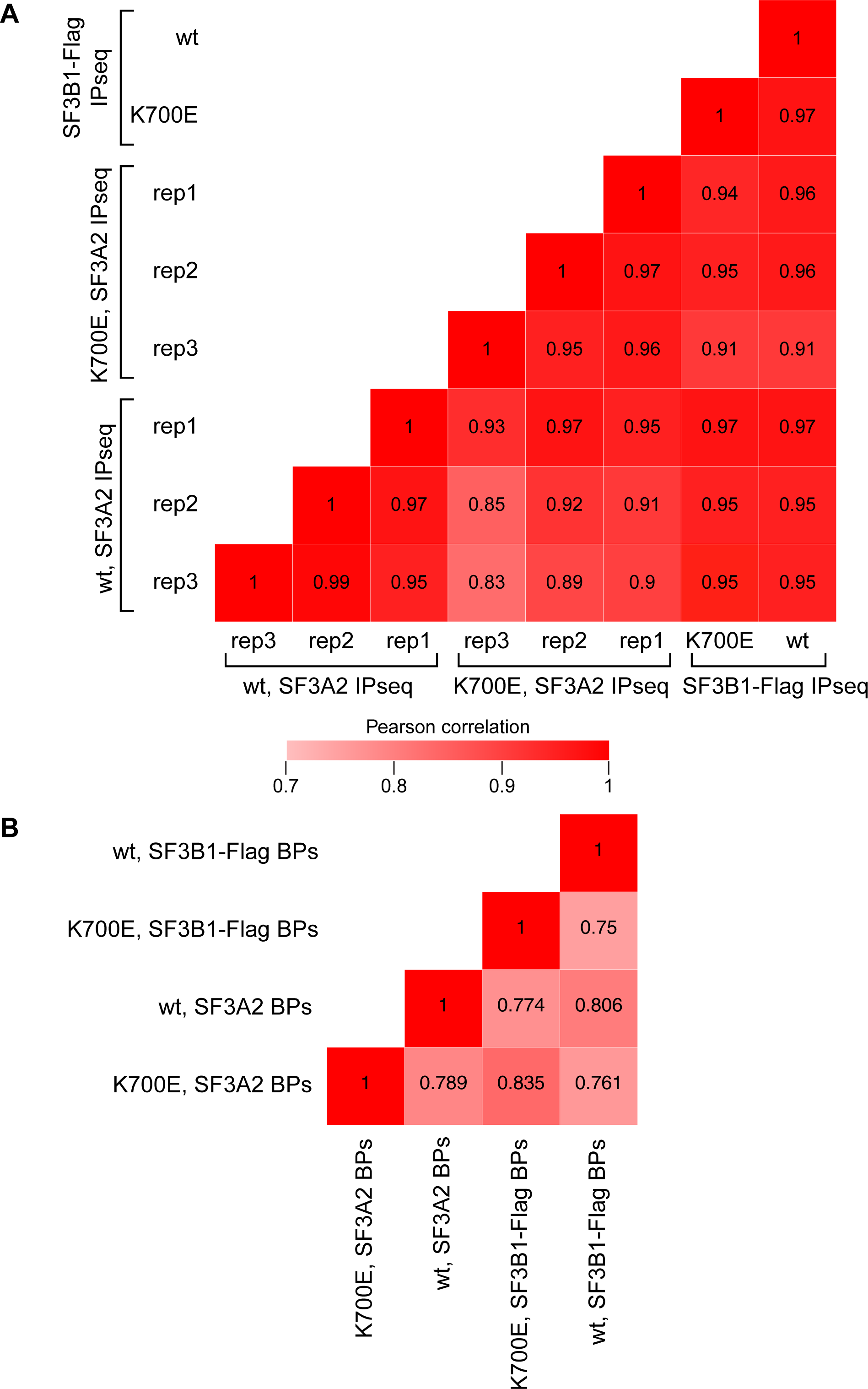
Branch sites and predicted branchpoints broadly correlate between WT and K700E samples. **A.** Correlation between IPseq RPKMs in 3’ ss and intronic clusters. **B.** Correlation of predicted BP coincidence. BPs ranked 1 in in 3’ss clusters were analyzed. (Related to Figure 2).

**Figure S6.**
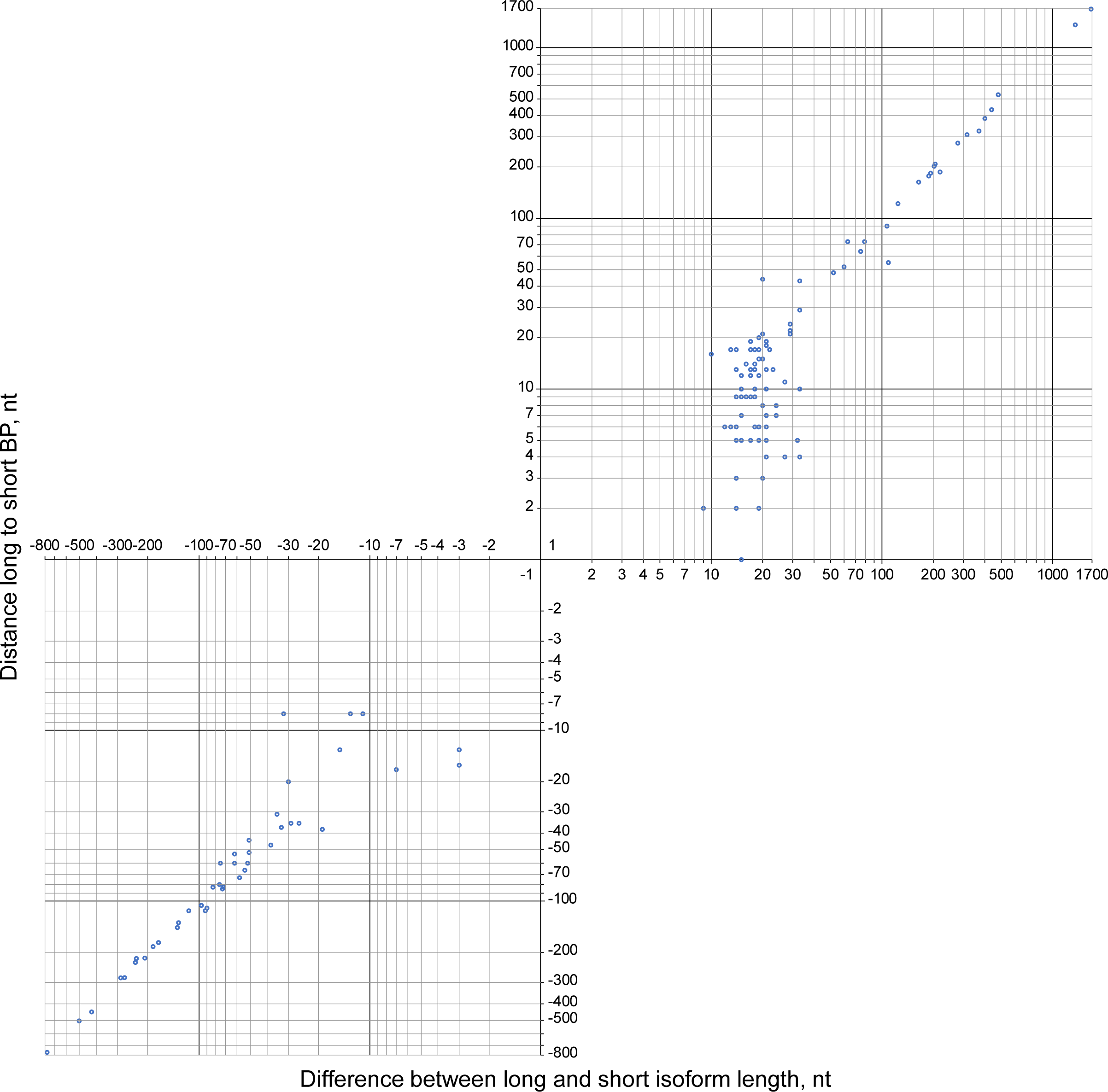
The distance between 3’ ss used in WT and K700E correlates with that between BPs selected in these cells (Related to Figure 3). Isoform length differences (the distances between selectively used 3’ ss) are plotted against the distances between WT and K700E BPs. Positive numbers indicate mutant 3’ss and BPs located upstream of WT; negative numbers indicate mutant 3’ss and BPs located downstream of WT.

**Figure S7.**
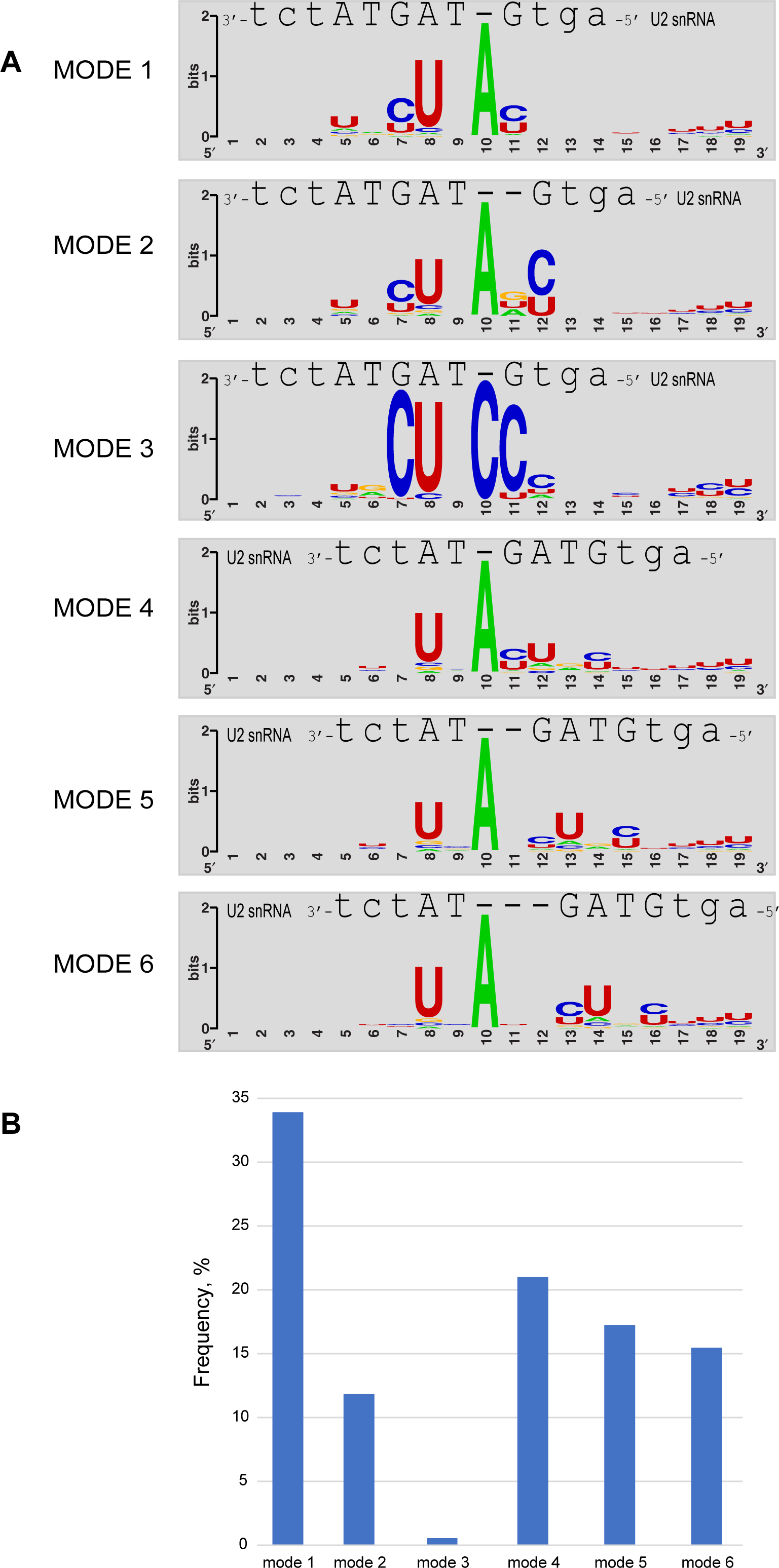
K562 branch sites base-pairing in different modes with U2 snRNA (Related to Figure 2). Pre-mRNA sequences flanking intronic branchpoints discovered in two studies of K562 cells (28, 32) were assessed for optimal base-pairing with the branch-site recognition sequence of U2 snRNA, GUAGUA. Consensus sequences of branch sites interacting with U2 in distinct base-pairing modes (29) were generated with WebLogo (49). **A.** The U2 GUAGUA branch interaction motif and its flanking sequence is shown in register above each consensus. **B.** The frequency of each described base-pairing mode from A.

**Figure S8.**
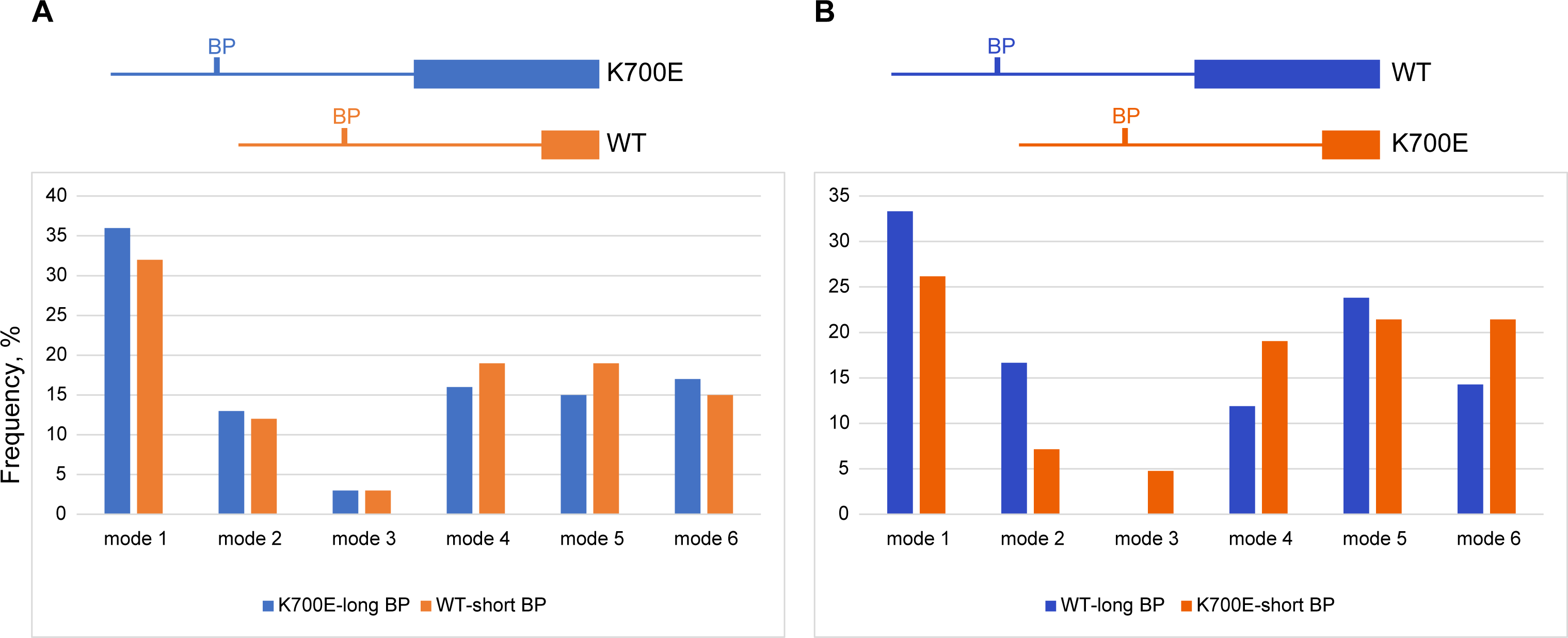
Base-pairing modes of differentially used branch sites in introns with alternative 3’ splice sites activated in SF3B1 K700E mutant cells. The frequency (%) of each described base-pairing mode at sites associated with long isoforms (**A**) and short isoforms (**B**) in K562 SF3B1-K700E cells is shown as in Figure S6B. The relative positions of alternative 3’ splice sites and their associated branchpoints are diagrammed above each panel.

**Figure S9.**
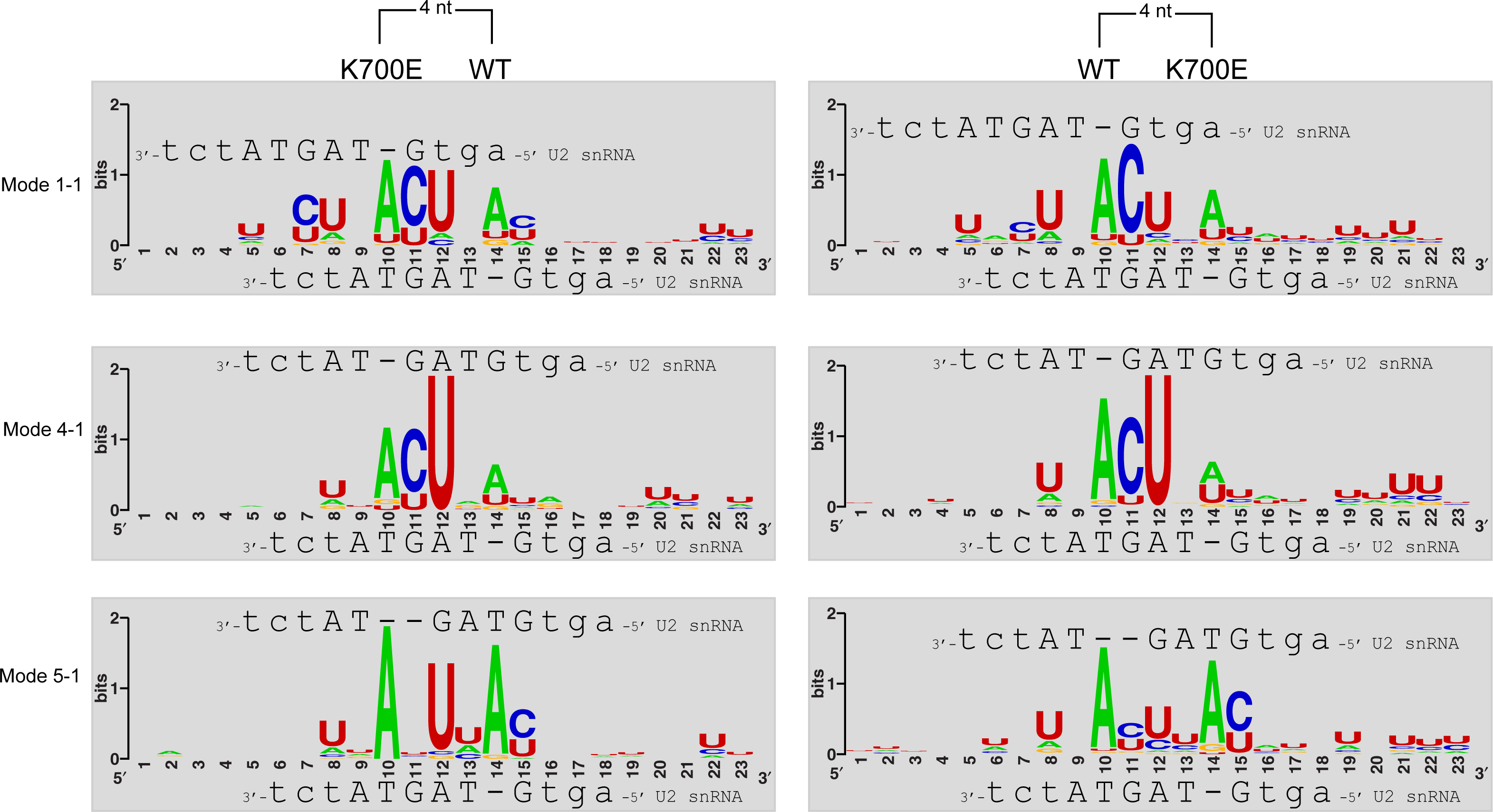
Consensus sequences of K700E preferred branch sites located 4 nt upstream or downstream of sites used in WT cells (Related to Figure 4). Mutant and WT preferred branch site pairs separated by 4 nt were grouped by U2 pairing modes. Consensus sequences are shown as in Figure S6, with the U2 sequence aligned to the upstream site above and the downstream site below. Only the most frequently occurring mode combinations are shown as indicated on the left.

**Figure S10.**
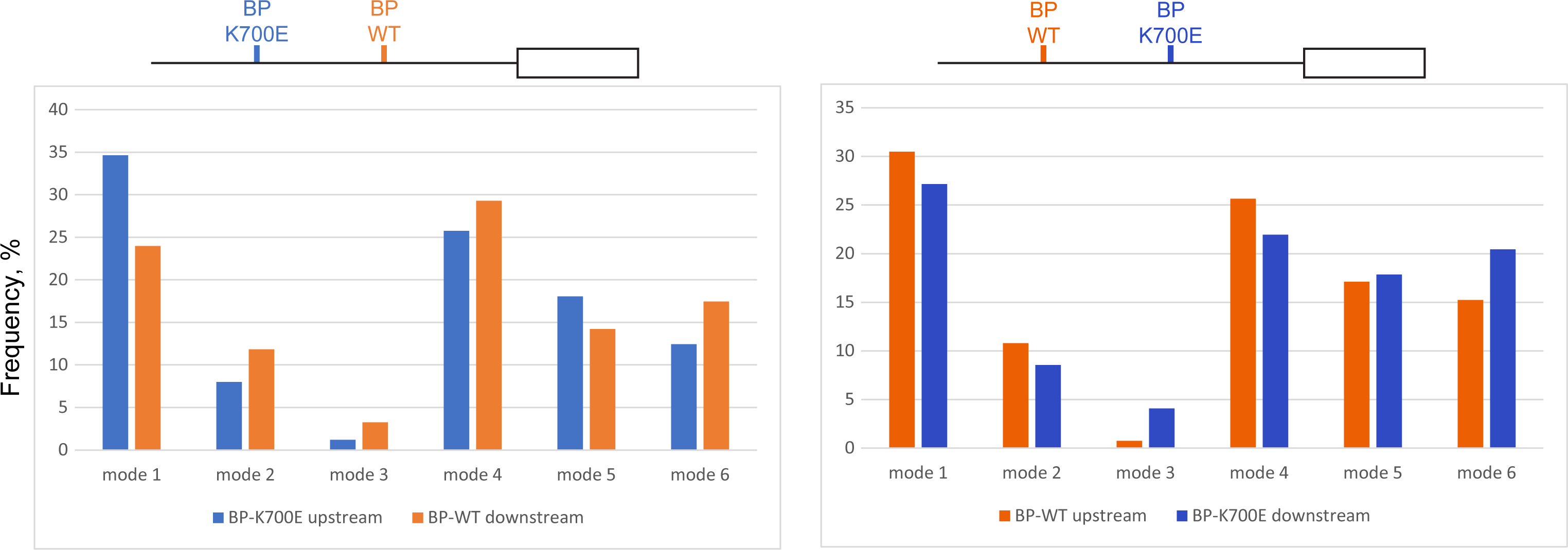
Base-pairing modes of K700E preferred and WT preferred branch site pairs separated by 7 to 14nt within branch site read clusters (Related to Figure 4). The frequency of each described base-pairing mode at preferentially used sites is shown as in Figure S6B.

**Figure S11.**
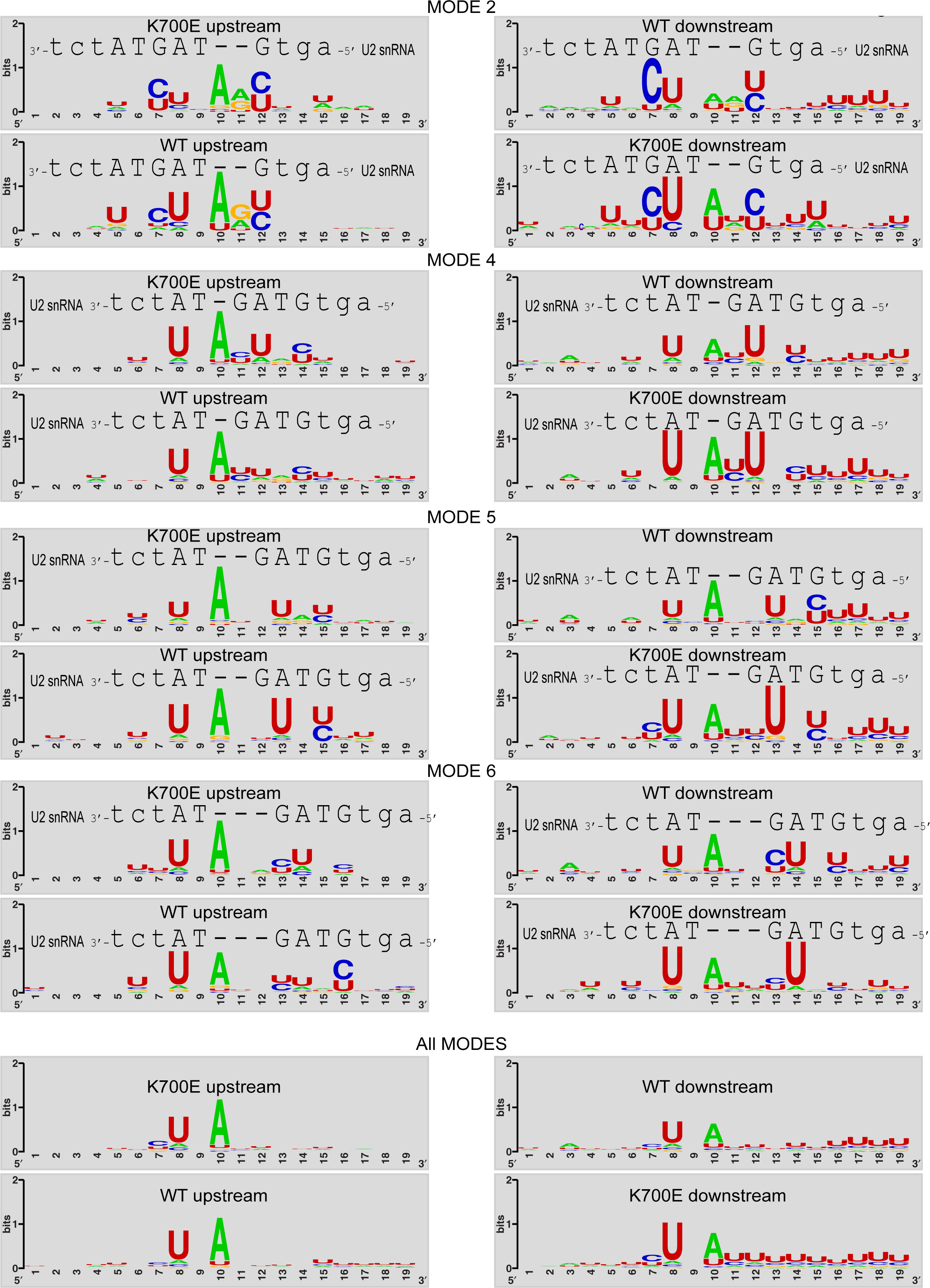
Consensus sequences of differentially used branch sites separated by 7 to 14 nt and base-pairing with U2 snRNA in non-canonical modes 2-6 are shown as in Figure 5. Consensus sequences from the combination of all these sites without their separation by base-pairing mode are shown at the bottom. (Related to Figure 5).

## References

1. K. Yoshida, et al., Frequent pathway mutations of splicing machinery in myelodysplasia. Nature 478, 64–69 (2011).

2. R. K. Bradley, O. Anczuków, RNA splicing dysregulation and the hallmarks of cancer. Nat Rev Cancer 23, 135–155 (2023).

3. Q. Zhang, Y. Ai, O. Abdel-Wahab, Molecular impact of mutations in RNA splicing factors in cancer. Molecular Cell 84, 3667–3680 (2024).

4. S. C.-W. Lee, et al., Synthetic Lethal and Convergent Biological Effects of Cancer-Associated Spliceosomal Gene Mutations. Cancer cell 34, 225 (2018).

5. L. Malcovati, et al., SF3B1 mutation identifies a distinct subset of myelodysplastic syndrome with ring sideroblasts. Blood 126, 233–241 (2015).

6. V. Quesada, et al., Exome sequencing identifies recurrent mutations of the splicing factor SF3B1 gene in chronic lymphocytic leukemia. Nat Genet 44, 47–52 (2012).

7. S. Alsafadi, et al., Cancer-associated SF3B1 mutations affect alternative splicing by promoting alternative branchpoint usage. Nat Commun 7, 10615 (2016).

8. S. Huber, et al., SF3B1 mutated MDS: Blast count, genetic co-abnormalities and their impact on classification and prognosis. Leukemia 36, 2894–2902 (2022).

9. R. B. Darman, et al., Cancer-Associated SF3B1 Hotspot Mutations Induce Cryptic 3′ Splice Site Selection through Use of a Different Branch Point. Cell Reports 13, 1033–1045 (2015).

10. J. Zhang, et al., Disease-Causing Mutations in SF3B1 Alter Splicing by Disrupting Interaction with SUGP1. Mol Cell 76, 82–95.e7 (2019).

11. C. DeBoever, et al., Transcriptome Sequencing Reveals Potential Mechanism of Cryptic 3’ Splice Site Selection in SF3B1-mutated Cancers. PLOS Computational Biology 11, e1004105 (2015).

12. M. E. Wilkinson, C. Charenton, K. Nagai, RNA Splicing by the Spliceosome. Annual Review of Biochemistry 89, 359–388 (2020).

13. C. van der Feltz, A. A. Hoskins, Structural and Functional Modularity of the U2 snRNP in pre-mRNA Splicing. Crit Rev Biochem Mol Biol 54, 443–465 (2019).

14. Z. Zhang, et al., Structural insights into how Prp5 proofreads the pre-mRNA branch site. Nature 596, 296–300 (2021).

15. Z. Zhang, et al., Structural insights into the cross-exon to cross-intron spliceosome switch. Nature 630, 1012–1019 (2024).

16. O. Gozani, J. Potashkin, R. Reed, A Potential Role for U2AF-SAP 155 Interactions in Recruiting U2 snRNP to the Branch Site. Mol Cell Biol 18, 4752–4760 (1998).

17. Z. Liu, et al., Pan-cancer analysis identifies mutations in SUGP1 that recapitulate mutant SF3B1 splicing dysregulation. Proceedings of the National Academy of Sciences 117, 10305–10312 (2020).

18. S. Alsafadi, et al., Genetic alterations of SUGP1 mimic mutant-SF3B1 splice pattern in lung adenocarcinoma and other cancers. Oncogene 40, 85–96 (2021).

19. J. Zhang, et al., DHX15 is involved in SUGP1-mediated RNA missplicing by mutant SF3B1 in cancer. Proc Natl Acad Sci U S A 119, e2216712119 (2022).

20. I. Beusch, et al., Targeted high-throughput mutagenesis of the human spliceosome reveals its in vivo operating principles. Molecular Cell 83, 2578–2594.e9 (2023).

21. H. M. Maul-Newby, et al., A model for DHX15 mediated disassembly of A-complex spliceosomes. RNA 28, 583–595 (2022).

22. Q. Feng, K. Krick, J. Chu, C. B. Burge, Splicing quality control mediated by DHX15 and its G-patch activator SUGP1. Cell Reports 42 (2023).

23. J. Zhang, et al., Characterization of the SF3B1–SUGP1 interface reveals how numerous cancer mutations cause mRNA missplicing. Genes Dev 37, 968–983 (2023).

24. S. Benbarche, et al., GPATCH8 modulates mutant SF3B1 mis-splicing and pathogenicity in hematologic malignancies. Molecular Cell 84, 1886–1903.e10 (2024).

25. Q. Tang, et al., SF3B1/Hsh155 HEAT motif mutations affect interaction with the spliceosomal ATPase Prp5, resulting in altered branch site selectivity in pre-mRNA splicing. Genes Dev. 30, 2710–2723 (2016).

26. B. Zhao, et al., Cancer-associated mutations in SF3B1 disrupt the interaction between SF3B1 and DDX42. The Journal of Biochemistry 172, 117–126 (2022).

27. F. Yang, et al., Mechanisms of the RNA helicases DDX42 and DDX46 in human U2 snRNP assembly. Nat Commun 14, 897 (2023).

28. T. R. Mercer, et al., Genome-wide discovery of human splicing branchpoints. Genome Res 25, 290–303 (2015).

29. A. J. Taggart, et al., Large-scale analysis of branchpoint usage across species and cell lines. Genome Res 27, 639–649 (2017).

30. B. Signal, B. S. Gloss, M. E. Dinger, T. R. Mercer, Machine learning annotation of human branchpoints. Bioinformatics 34, 920–927 (2018).

31. P. Zhang, et al., Genome-wide detection of human variants that disrupt intronic branchpoints. Proc Natl Acad Sci U S A 119, e2211194119 (2022).

32. Y. Zeng, et al., Profiling lariat intermediates reveals genetic determinants of early and late co-transcriptional splicing. Mol Cell 82, 4681–4699.e8 (2022).

33. A. Damianov, et al., The splicing regulators RBM5 and RBM10 are subunits of the U2 snRNP engaged with intron branch sites on chromatin. Molecular Cell 84, 1496–1511.e7 (2024).

34. Z. Zhang, et al., Molecular architecture of the human 17S U2 snRNP. Nature 583, 310–313 (2020).

35. Y. K. Lieu, et al., SF3B1 mutant-induced missplicing of MAP3K7 causes anemia in myelodysplastic syndromes. Proceedings of the National Academy of Sciences 119, e2111703119 (2022).

36. S. Shen, et al., rMATS: robust and flexible detection of differential alternative splicing from replicate RNA-Seq data. Proc Natl Acad Sci U S A 111, E5593–5601 (2014).

37. C. W. J. Smith, E. B. Porro, J. G. Patton, B. Nadal-Ginard, Scanning from an independently specified branch point defines the 3′ splice site of mammalian introns. Nature 342, 243–247 (1989).

38. C. W. J. Smith, T. T. Chu, B. Nadal-Ginard, Scanning and Competition between AGs Are Involved in 3’ Splice Site Selection in Mammalian Introns. Molecular and Cellular Biology 13, 4939–4952 (1993).

39. R. Yoshimoto, N. Kataoka, K. Okawa, M. Ohno, Isolation and characterization of post-splicing lariat-intron complexes. Nucleic Acids Res 37, 891–902 (2009).

40. J.-B. Fourmann, et al., The target of the DEAH-box NTP triphosphatase Prp43 in Saccharomyces cerevisiae spliceosomes is the U2 snRNP-intron interaction. eLife 5, e15564 (2016).

41. R. Toroney, K. H. Nielsen, J. P. Staley, Termination of pre-mRNA splicing requires that the ATPase and RNA unwindase Prp43p acts on the catalytic snRNA U6. Genes Dev. 33, 1555–1574 (2019).

42. J. C. S. Noble, Z.-Q. Pan, C. Prives, J. L. Manley, Splicing of SV40 early pre-mRNA to large T and small t mRNAs utilizes different patterns of lariat branch sites. Cell 50, 227–236 (1987).

43. R. A. Padgett, et al., Nonconsensus branch-site sequences in the in vitro splicing of transcripts of mutant rabbit beta-globin genes. Proc Natl Acad Sci U S A 82, 8349– 8353 (1985).

44. B. Ruskin, J. M. Greene, M. R. Green, Cryptic branch point activation allows accurate in vitro splicing of human β-globin intron mutants. Cell 41, 833–844 (1985).

45. E. Glasser, et al., Pre-mRNA splicing factor U2AF2 recognizes distinct conformations of nucleotide variants at the center of the pre-mRNA splice site signal. Nucleic Acids Res 50, 5299–5312 (2022).

46. M. Corioni, N. Antih, G. Tanackovic, M. Zavolan, A. Krämer, Analysis of in situ pre-mRNA targets of human splicing factor SF1 reveals a function in alternative splicing. Nucleic Acids Res 39, 1868–1879 (2011).

47. S. Sharma, S. P. Wongpalee, A. Vashisht, J. A. Wohlschlegel, D. L. Black, Stem– loop 4 of U1 snRNA is essential for splicing and interacts with the U2 snRNP-specific SF3A1 protein during spliceosome assembly. Genes Dev 28, 2518–2531 (2014).

48. A. J. Taggart, et al., Large-scale analysis of branchpoint usage across species and cell lines. Genome Res 27, 639–649 (2017).

49. G. E. Crooks, G. Hon, J.-M. Chandonia, S. E. Brenner, WebLogo: A Sequence Logo Generator. Genome Res 14, 1188–1190 (2004).

